# Targeting advanced prostate cancer with STEAP1 chimeric antigen receptor T cell therapy

**DOI:** 10.1101/2022.05.16.492156

**Authors:** Vipul Bhatia, Nikhil V. Kamat, Tiffany E. Pariva, Li-Ting Wu, Annabelle Tsao, Koichi Sasaki, Lauren T. Wiest, Ailin Zhang, Dmytro Rudoy, Roman Gulati, Radhika A. Patel, Martine P. Roudier, Lawrence D. True, Michael C. Haffner, Peter S. Nelson, Saul J. Priceman, Jun Ishihara, John K. Lee

## Abstract

Six transmembrane epithelial antigen of the prostate 1 (STEAP1) is a compelling tumor-associated cell surface antigen for therapeutic targeting in solid tumors. We identified broad expression of STEAP1 (87% positive) in lethal metastatic prostate cancer, even more so than prostate-specific membrane antigen (PSMA, 60% positive) which is a clinically established diagnostic and therapeutic target. Second-generation chimeric antigen receptor (CAR) T cells were engineered for reactivity against STEAP1 and demonstrated substantial antitumor activity in metastatic human prostate cancer models in immunodeficient mice. Adoptive transfer of STEAP1 CAR T cells was associated with prolonged peripheral persistence and either disease eradication or substantial tumor growth inhibition with progressive disease demonstrating antigen loss. As STEAP1 CAR T cells were also highly active in antigen density conditions as low as ∼1,500 molecules/cell, we generated a human STEAP1 (hSTEAP1) knock-in (KI) mouse to evaluate the potential for on-target off-tumor toxicities. hSTEAP1-KI mice demonstrated a pattern of systemic hSTEAP1 expression akin to that observed in humans with the greatest expression found in the prostate gland. Mouse-in-mouse studies of STEAP1 CAR T cell therapy in immunocompetent hSTEAP1-KI mice engrafted with disseminated mouse prostate cancer showed preliminary safety without evidence of gross toxicity, cytokine storm, or architectural disruption and increased T cell infiltration at sites of systemic hSTEAP1 expression. Tumor responses and extension of survival were appreciated but antigen loss was identified in recurrent and progressive disease. In summary, we report the extent of STEAP1 expression in treatment-refractory metastatic prostate cancer, the generation of a STEAP1 CAR T cell therapy with promising potency and safety in preclinical studies of advanced prostate cancer, and antigen escape as a mechanism of resistance to effective STEAP1 CAR T cell therapy.

## Introduction

Metastatic prostate cancer represents an incurable disease responsible for over 33,000 deaths per year in the United States^1^. Prostate cancer is critically reliant on androgen receptor (AR) signaling and thus the suppression of gonadal androgen production through surgical or chemical castration (androgen deprivation therapy) has been a mainstay of treatment for advanced disease. However, metastatic prostate cancer inevitably develops resistance to androgen deprivation therapy and enters a stage called metastatic castration-resistant prostate cancer (mCRPC). mCRPC is currently incurable and is considered the end-stage of the disease and is associated with a median overall survival of three years^2^. In the past decade, multiple therapies including an inhibitor of extragonadal androgen synthesis (abiraterone acetate)^3^, second-generation AR antagonists (enzalutamide)^4^, radioactive isotope (radium-223)^5^, and a prostate-specific membrane antigen (PSMA)-specific radioligand therapy (lutetium Lu 177 vipivotide tetraxetan)^6^ have been approved for mCRPC. Each of these agents extends survival on average by several months but long-term remissions are rare.

Strategies to reprogram the immune system to combat prostate cancer first gained traction with the clinical approval of the dendritic cell vaccine sipuleucel-T for asymptomatic mCRPC^7^. More recently, several types of immunotherapies including immune checkpoint inhibitors, a DNA cancer vaccine, antibody-drug conjugates (ADC), T cell engaging bispecific antibodies (T-BsAb), and chimeric antigen receptor (CAR) T cell therapies have been under active clinical investigation. CARs are synthetic receptors that leverage the potency, expansion, and memory of T cells and can be engineered against virtually any tumor-associated cell surface antigen.

The adoptive transfer of CAR T cells has rapidly become an established treatment for hematologic malignancies with exceptional response rates leading to six clinical approvals in the last five years. In contrast, CAR T cell therapies targeting solid tumors have lagged due to additional challenges related to the lack of *bona fide* tumor-specific antigens, inhospitable tumor microenvironments, and poor trafficking, persistence, and expansion of CAR T cells.

Despite the challenges observed in driving effective immune responses toward solid tumors, recent early phase clinic trials investigating CAR T cell therapies targeting PSMA in mCRPC have reported safety and evidence of significant biochemical and radiographic responses^8, 9^. These preliminary results serve to embolden efforts to develop and optimize new CAR T cell therapies for prostate cancer. While PSMA is the preeminent target for therapeutic and diagnostic development in prostate cancer, recent work indicates that PSMA expression may be quite heterogeneous in mCRPC^10^. Tumor antigen heterogeneity, especially in the context of single antigen-targeted CAR T cell therapies for solid tumors like prostate cancer, is an important barrier to therapeutic efficacy^11^. Thus, identifying cell surface antigens with broad and relatively homogeneous expression in prostate cancer is imperative. In addition, very few if any tumor-associated antigens demonstrate tumor-restricted expression—most also exhibit low level expression in normal tissues that could represent liabilities for CAR T cell therapies due to on-target off-tumor toxicities which can lead to devastating consequences including death^12^.

We previously performed integrated transcriptomic and cell surface proteomic profiling of human prostate adenocarcinoma cell lines and identified six transmembrane epithelial antigen of the prostate 1 (STEAP1) as one of the most highly enriched cell surface antigens^13^. STEAP1 was first described over two decades ago^14^ and was recognized as being highly expressed in prostate cancer. STEAP1 is strongly expressed in >80% of mCRPC with bone or lymph node involvement^15^, 62% of Ewing sarcoma^16^, and multiple other cancer types^17^. STEAP1 belongs to the STEAP family of metalloreductases that can form homotrimers or heterotrimers with other STEAP proteins^18^. STEAP1 has an established functional role in promoting cancer cell proliferation, invasion, and epithelial-to-mesenchymal transition^19–23^. Furthermore, STEAP1 demonstrates limited expression in normal tissue^24^ which makes it a highly compelling target for cancer therapy.

Multiple immunotherapeutic agents have been developed to target STEAP1 in cancer but an approach employing CAR T cell therapy has not yet been reported. The ADC vandortuzumab vedotin (DSTP3086S) consisting of a humanized anti-STEAP1 IgG1 antibody linked to monomethyl auristatin E was found to have an acceptable safety profile in a phase I clinical trial in mCRPC but few objective tumor responses were observed^25^. A T-BsAb incorporating two anti-STEAP1 fragment-antigen binding (Fab) domains, an anti-CD3 single chain variable fragment (scFv), and a fragment crystallizable (Fc) domain engineered to lack effector function called AMG 509 is currently being evaluated in a phase I clinical trial in mCRPC^26^. A symmetric dual bivalent T-BsAb called BC261 was also recently reported to demonstrate potent antitumor activity across multiple preclinical models of prostate cancer and Ewing sarcoma^27^. In addition, a human leukocyte antigen (HLA) class I-restricted T cell receptor (TCR) specific for a STEAP1 peptide has been shown to inhibit local and metastatic Ewing sarcoma growth in a preclinical xenograft model after adoptive transfer of transgenic T cells^28^.

In this study, we performed comparative analysis of the relative expression of STEAP1 and PSMA in lethal mCRPC to investigate the utility of targeting STEAP1 in the current era of PSMA theranostics. We engineered and screened second-generation STEAP1 CARs for antigen-specific T cell activation and target cell cytolysis which yielded a lead candidate for further characterization. We determined the functional epitope specificity of STEAP1 CAR T cells and profiled the expansion and immunophenotype of STEAP1 CAR T cell products from multiple donors. We then established the potency and preliminary safety of STEAP1 CAR T cell therapy in relevant preclinical human-in-mouse and mouse-in-mouse models of prostate cancer. Collectively, these studies provide strong rationale for the clinical translation of STEAP1 CAR T cell therapy to men with mCRPC and guide future studies to overcome potential mechanisms of therapeutic resistance.

## Results

### STEAP1 is broadly expressed in treatment-refractory mCRPC tissues

We first set out to determine the pattern and extent of STEAP1 expression relative to PSMA in advanced metastatic prostate cancer. We performed immunohistochemical (IHC) staining on a duplicate set of tissue microarrays consisting of 121 metastatic tumors (each with up to three cores represented) collected from 45 men with lethal mCRPC patients collected by rapid autopsy between the years 2010 and 2017 through the University of Washington Tumor Acquisition Necropsy Program^29^ (**figure 1A**). Plasma membrane staining for STEAP1 and PSMA in each tissue was scored by a research pathologist and semiquantitative H-scores were determined based on the staining intensity (0, 1, 2, or 3, **supplemental figure 1A**) multiplied by the percentage of cancer cells staining at each intensity (**figure 1B**). Based on these results, we used a generalized linear mixed statistical model to determine that the odds of non-zero staining was 7.7-fold (95% CI 2.8 to 20.8, p<0.001) higher for STEAP1 than for PSMA. By implementing a minimal staining threshold with an H-score cut-off of 30, we found that 87.7% of evaluable matched mCRPC tissues (100 of 114) demonstrated staining for STEAP1 compared to only 60.5% (69 of 114) for PSMA (**figure 1C**). In addition, 28.1% of mCRPC tissues (32 of 114) showed STEAP1 but not PSMA staining (**figure 1D**) whereas only 0.9% (one of 114) exhibited PSMA but not STEAP1 staining. We also observed several cases with heterogeneous expression of PSMA within cores (**figure 1E**) which is consistent with a recent report of intratumoral PSMA heterogeneity in mCRPC biopsies^10^.

**Figure 1.**
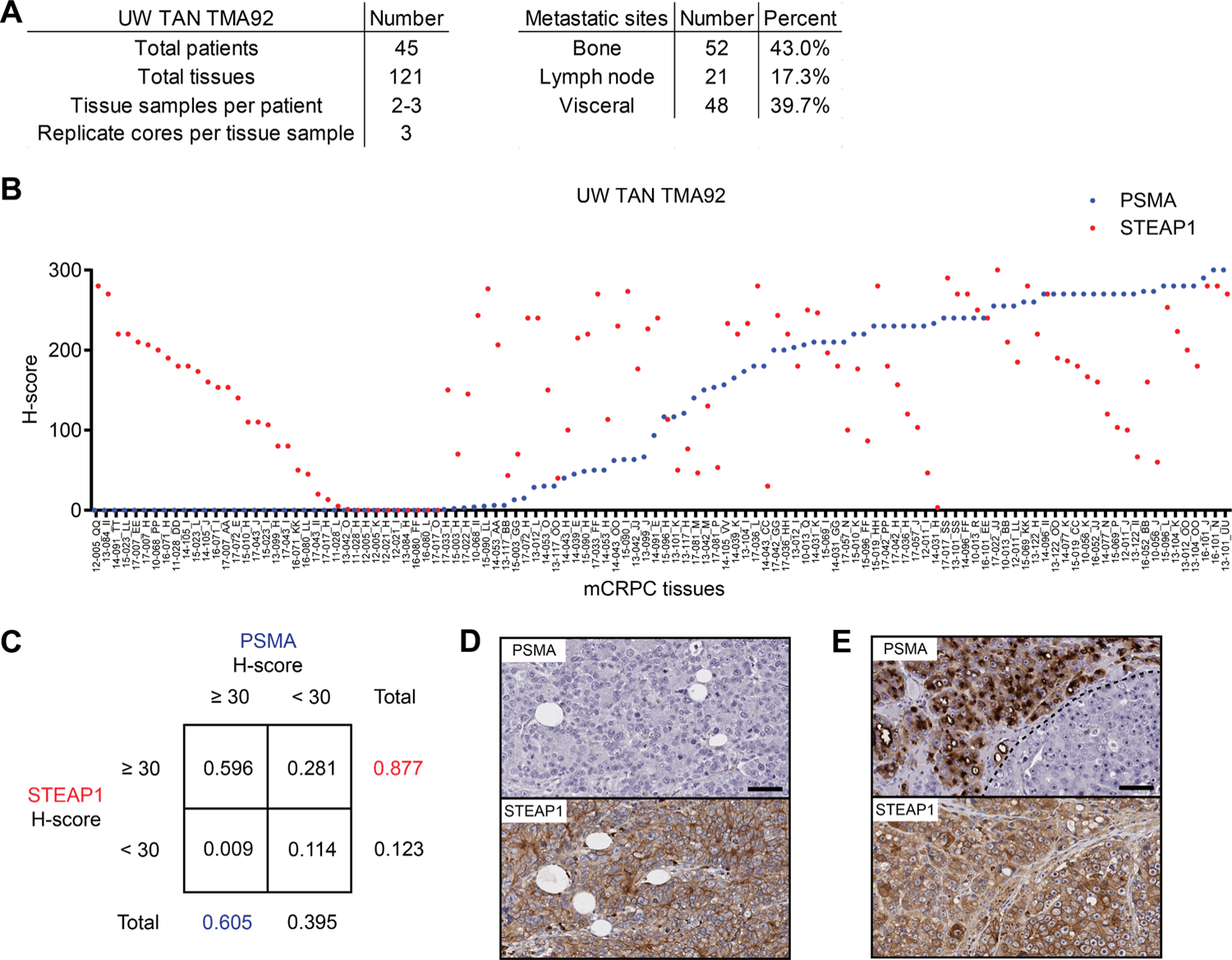
Comparative analysis of STEAP1 and PSMA in lethal, metastatic castration-resistant prostate cancer (mCRPC). (**A**) Characteristics of the mCRPC tissues represented on University of Washington Tissue Acquisition Necropsy Tissue Microarray 92 (UW TAN TMA92). (**B**) Plot showing paired average H-scores of STEAP1 (red) and PSMA (blue) immunohistochemical (IHC) staining of all cores from each mCRPC tissue. (**C**) Contingency table showing the frequency of mCRPC tissues with STEAP1 or PSMA IHC staining above or below an H-score threshold of 30. Micrographs of select mCRPC tissues after STEAP1 and PSMA IHC staining to highlight the (**D**) absence of PSMA but presence of STEAP1 expression and (**E**) intratumoral heterogeneity of PSMA expression but not STEAP1. Scale bars = 50 µm.

STEAP1 staining based on the minimal staining threshold was identified in 96% (48 of 50) of bone metastases, 95% (19 of 20) of lymph node metastases, and 76.6% (36 of 47) of visceral metastases (**supplemental figure 1B**). No difference in STEAP1 staining intensity was observed between bone and lymph node or lymph node and visceral metastatic sites. However, bone metastases demonstrated a higher STEAP1 H-score than visceral metastases (183.6 vs. 121.9, p=0.0018). We identified a positive Pearson correlation (r=0.3057, 95% CI 0.1314 to 0.4616, p<0.001) between the expression of STEAP1 and AR in cases represented on the tissue microarray (**supplemental figure 1C**) which was expected given that *STEAP1* is an androgen-regulated gene^30, 31^. In contrast, a negative correlation (r=-0.2172, 95% CI −0.3843 to −0.03628, p=0.0192) was appreciated between the expression of STEAP1 and the neuroendocrine differentiation marker synaptophysin (**supplemental figure 1D**). These findings suggest that, like PSMA^32^, STEAP1 expression may be lost with neuroendocrine transdifferentiation of prostate cancer.

### Development of a potent, antigen-specific STEAP1 CAR

Given the widespread expression of STEAP1 in late-stage mCRPC and its reported functional role in cancer progression^24, 33, 34^, we next started to engineer a lentiviral STEAP1-specific second-generation CAR. We used the pCCL-c-MNDU3-X lentiviral backbone^35^ which has been widely used for hematopoietic stem cell gene therapy^36^ and CAR expression in T cells driven by the internal MNDU3 promoter has been shown to be higher than that achieved with an EFS promoter^37^. A 4-1BB costimulatory domain was favored due to its association with T cell memory formation and prolonged persistence^38^ and a CD28 transmembrane domain was introduced as this has been shown to reduce the antigen threshold for second-generation 4-1BB CAR T cell activation^39^. We incorporated the fully humanized scFv derived from vandortuzumab vedotin, an ADC targeting STEAP1 whose development was discontinued after a phase I clinical trial^25^. This scFv is a humanized variant of the murine monoclonal antibody (mAb 120.545) originally developed by Agensys, Inc. that demonstrates 1 nM affinity in cell-based binding assays^40^. To potentially tune CAR activity, we implemented three different hinge/spacer lengths including short (IgG4 hinge), medium (IgG4 hinge-CH3), and long (IgG4 hinge-CH2-CH3). The long spacer was engineered with previously described 4/2-NQ mutations^41^ in the CH2 domain to prevent Fc-gamma receptor binding and activation-induced cell death that occurs with the adoptive transfer of long spacer CAR T cells into immunodeficient mice. The three candidate CARs were cloned into the lentiviral vector (**figure 2A**) that also co-expresses truncated epidermal growth factor receptor (EGFRt) as a transduction marker. Lentiviruses were generated and used to transduce human CD4 and CD8 T cells enriched from human donor peripheral blood mononuclear cells (PBMCs) collected from pheresis. Expanded CD4 and CD8 CAR T cells were immunophenotyped (**supplemental figure 2A**) and reconstituted into cell products of a defined composition with a normal CD4/CD8 ratio to evaluate their functional activities.

**Figure 2.**
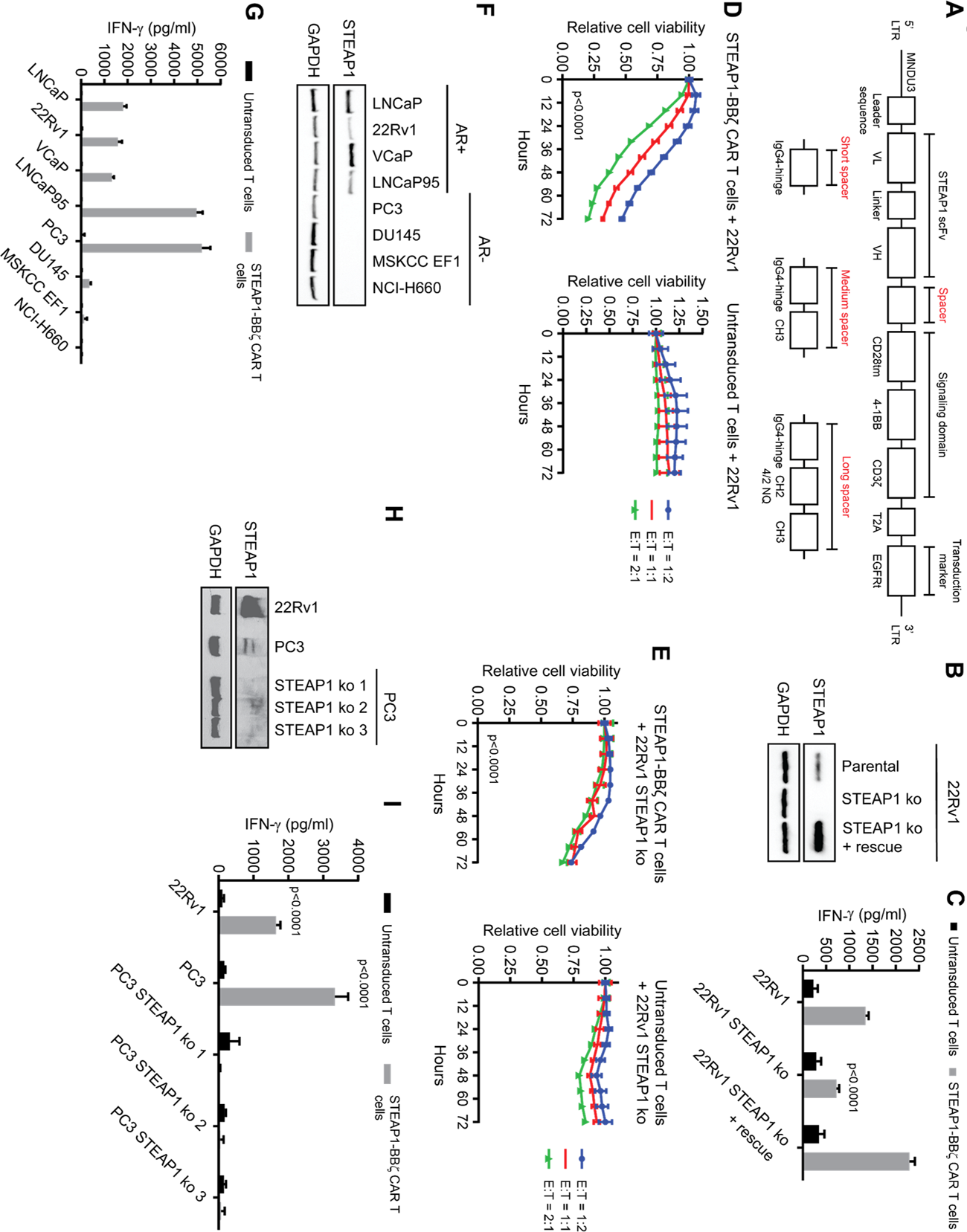
Screening second-generation 4-1BB chimeric antigen receptors (CARs) to identify a lead for STEAP1 CAR T cell therapy. (**A**) Schematic of the lentiviral STEAP1 CAR construct and variation based on short, medium, and long spacers. LTR = long terminal repeat; MNDU3 = Moloney murine leukemia virus U3 region; scFv = single-chain variable fragment; VL = variable light chain; VH = variable heavy chain; tm = transmembrane; EGFRt = truncated epidermal growth factor receptor; 4/2 NQ = CH2 domain mutations to prevent binding to Fc-gamma receptors. (**B**) Immunoblot analysis showing expression of STEAP1 in 22Rv1 parental cells, 22Rv1 STEAP1 knockout (ko) cells, and 22Rv1 STEAP1 ko cells with rescue of STEAP1 expression by lentiviral expression. GAPDH is used as a protein loading control. (**C**) IFN-γ enzyme-linked immunosorbent assay (ELISA) results from co-cultures of either untransduced T cells or STEAP1-BBζ CAR T cells with each of the 22Rv1 sublines at a 1:1 ratio at 24 hours. n = 4 replicates per condition. Bars represent SD. (**D**) Relative cell viability of 22Rv1 target cells over time measured by fluorescence live cell imaging upon co-culture with (left) STEAP1-BBζ CAR T cells or (right) untransduced T cells at variable effector-to-target (E:T) cell ratios. (**E**) Relative cell viability of 22Rv1 STEAP1 ko target cells over time measured by fluorescence live cell imaging upon co-culture with (left) STEAP1-BBζ CAR T cells or (right) untransduced T cells at variable E:T cell ratios. In **D** and **E**, n = 4 replicates per condition and bars represent SEM. (**F**) Immunoblot analysis demonstrating expression of STEAP1 in androgen receptor (AR)-positive human prostate cancer cell lines but not AR-negative prostate cancer cell lines. GAPDH is used as a protein loading control. (**G**) IFN-γ quantification by ELISA from co-cultures of either untransduced T cells or STEAP1-BBζ CAR T cells with each of the human prostate cancer cell lines in **F** at a 1:1 ratio at 24 hours. n = 4 replicates per condition. Bars represent SD. (**H**) Immunoblot analysis with prolonged exposure to evaluate expression of STEAP1 in 22Rv1, PC3, and PC3 STEAP1 ko sublines. GAPDH is used as a protein loading control. (**I**) IFN-γ quantification by ELISA from co-cultures of either untransduced T cells or STEAP1-BBζ CAR T cells with each of the human prostate cancer cell lines in **F** at a 1:1 ratio at 24 hours. n = 4 replicates per condition. Error bars represent SEM. For panel **C** and **I**, two-way ANOVA with Sidak’s multiple comparison test was used. For panels **D** and **E**, two-way ANOVA with Tukey’s multiple comparisons test was used.

To control for STEAP1 expression in an isogenic manner, we focused on the 22Rv1 human prostate cancer cell line that demonstrates native STEAP1 expression and performed STEAP1 knockout (ko) by CRISPR/Cas9 genome editing. We then generated a STEAP1 rescue line from the 22Rv1 STEAP1 ko by transduction with a STEAP1 expressing lentivirus (**figure 2B**). These lines were then used to screen the three short, medium, and long spacer STEAP1 CAR T cells in co-culture assays with a readout of interferon-gamma (IFN-γ) release as an indicator of T cell activation. Only the long spacer STEAP1 CAR T cells (hereafter called STEAP1-BBζ CAR T cells) demonstrated the anticipated antigen-specific pattern of IFN-γ release (**figure 2C, supplemental figure 2B**). Further, STEAP1-BBζ CAR T cells showed substantial dose-dependent cytolysis of 22Rv1 cells compared to untransduced T cells (**figure 2D**) and demonstrated relative sparing of 22Rv1 STEAP1 ko cells (**figure 2E**). Similar studies were then performed in the DU145 human prostate cancer cell line that lacks native STEAP1 expression but was engineered to express STEAP1 (DU145 STEAP1) by lentiviral transduction. In this setting, STEAP1-BBζ CAR T cell activation was only observed in co-cultures with DU145 STEAP1 cells and not the parental DU145 cells (**supplemental figure 2C**). Cytolytic activity was only appreciated with STEAP1-BBζ CAR T cells and not untransduced T cells in co-cultures with DU145 STEAP1 cells (**supplemental figure 2D**).

We subsequently analyzed a larger panel of human prostate cancer cell lines to characterize their native STEAP1 expression by immunoblot analysis. The cell lines with known AR expression/activity (LNCaP, 22Rv1, VCaP, and LNCaP95) showed varying levels of STEAP1 expression while the AR-null cell lines (PC3, DU145, MSKCC EF1, and NCI-H660) did not appear to express detectable levels of STEAP1 (**figure 2F**). We proceeded to perform co-cultures of STEAP1-BBζ CAR T with these lines to further validate their antigen-specific activation based on IFN-γ release (**figure 2G**). However, we observed a discordant finding in that the PC3 line, which showed no apparent STEAP1 protein expression (**figure 2F**), induced substantial activation of STEAP1-BBζ CAR T cells. Prior literature suggested that STEAP1 is expressed in the PC3 cell line at low levels^42^. Indeed, prolonged immunoblot exposure revealed a band suggesting the presence of very low expression of STEAP1 (**figure 2H**). To confirm whether the STEAP1-BBζ CAR T cell activation was due to this minor STEAP1 expression in PC3 cells, we generated three PC3 STEAP1 ko sublines (**figure 2H**) and again performed co-cultures with STEAP1-BBζ CAR T cells. STEAP1 ko in the PC3 line led to the abrogation of STEAP1-BBζ CAR T cell activation (**figure 2I**), further validating specificity and providing evidence of the sensitivity of STEAP1-BBζ CAR T cells to low antigen density conditions.

### Lack of cross-reactivity of STEAP1-BBζ CAR with mouse Steap1 and human STEAP1B

Consistent with the anti-human specificity of vandortuzumab vedotin, STEAP1*-*BBζ CAR T cells did not demonstrate cross reactivity with mouse Steap1 (**supplemental figure 3A-C**). However, we used this as an opportunity to individually reconstitute the three human STEAP1 extracellular domains (ECDs) onto mouse Steap1 (**supplemental figure 3D**) to determine which ECDs are critical for epitope recognition by STEAP1*-*BBζ CAR T cells. Co-culture experiments were performed with STEAP1-BBζ CAR T cells and DU145 cells engineered to express mouse Steap1 with individual replacement of mouse ECDs with human ECDs. We found that human STEAP1 ECD2 but not ECD1 or ECD3 was associated with STEAP1-BBζ CAR T cell activation (**supplemental figure 3E**). Interestingly, the human STEAP1 and mouse Steap1 ECD2 demonstrate 93.9% (31/33 amino acids) homology (**supplemental figure 3F**), indicating that Q198 and/or I209 of human STEAP1 are critical to productive recognition by STEAP1-BBζ CAR T cells. Q198 has been shown to interact with the Fab of 120.545 as part of an interaction hotspot based on a recent structure resolved by cryogenic electron microscropy^18^.

Of the human STEAP family of proteins, STEAP1B has the greatest homology to STEAP1^42^. Three STEAP1B transcripts have been identified, of which all demonstrate complete conservation of the amino acid sequence of human STEAP1 ECD2 (**supplemental figure 4A**). The consensus membrane topology prediction algorithm TOPCONS^43^ projected these sequences as being extracellular in the three STEAP1B protein isoforms (**supplemental figure 4B**) albeit with low reliability scores due to a lack of consensus between models (**supplemental figure 4C**). Prior analysis using a hidden Markov model had also suggested that this sequence could be intracellular rather than extracellular in STEAP1B protein isoforms 1 and 2^42^. However, the crystal structure of STEAP1B has not yet been determined to directly substantiate these predictions. To functionally evaluate whether STEAP1-BBζ CAR T cells might also be reactive against STEAP1B, we performed co-cultures using DU145 lines engineered to express each of the three isoforms of STEAP1B. We did not identify evidence of STEAP1-BBζ CAR T cell activation (**supplemental figure 4D**), suggesting that the STEAP1 epitope recognized by STEAP1-BBζ CAR T cells is not presented by STEAP1B despite apparent sequence homology.

### Characterization of STEAP1-BBζ CAR T cell products across a series of donors

We next profiled the expansion, transduction efficiency, and immunophenotype of STEAP1-BBζ CAR T cell products using three independent sets of peripheral blood mononuclear cells (PBMCs) collected from healthy donors. We generally observed a 20- to 40-fold expansion of STEAP1-BBζ CAR T cells within 11 days of culture (**supplementary figure 5A**). The percentage of EGFRt^+^ CD8 T cells ranged from 24.3% to 54.2% while the percentage of EGFRt^+^ CD4 T cells was higher and ranged from 60.1% to 74.9% in our STEAP1-BBζ CAR T cell products (**supplementary figure 5B**). We examined the expression of the T cell exhaustion markers PD-1 and LAG-3 in the untransduced and STEAP1-BBζ CAR T cell subsets and observed no significant increase in expression (**supplementary figure 5C**). This finding suggested low or absent tonic signaling by the STEAP1-BBζ CAR which was encouraging as constitutive CAR signaling can negatively impact CAR T cell effector function^44^.

Both stem cell memory T cell (Tscm) and central memory T cell (Tcm) phenotypes have been associated with the therapeutic efficacy of CAR T cell therapy as they promote sustained proliferation and persistence *in vivo*^45–47^. Immunophenotyping of untransduced and STEAP1-BBζ CAR T cell subsets demonstrated higher frequencies of Tscm cells compared to the T cell subsets in donor PBMCs from which the cell products were derived (**supplementary figure 5D**). This effect is likely due to the addition of IL-7 and/or IL-15 to the T cell expansion media as these cytokines have been shown to preserve and enhance Tscm differentiation^47, 48^. Our analysis also revealed an enrichment in Tcm populations particularly in the CD8 STEAP1-BBζ CAR T cells (**supplementary figure 5E**).

### STEAP1-BBζ CAR T cells demonstrate substantial antitumor effects in disseminated prostate cancer models with native STEAP1 expression established in immunodeficient mice

As an initial screen for *in vivo* antitumor activity, we established 22Rv1 subcutaneous xenograft tumors in male NOD scid gamma (NSG) mice. When tumors grew to approximately 100 mm^3^, mice were treated with a single intratumoral injection of either 5 x 10^6^ untransduced T cells or STEAP1-BBζ CAR T cells. Intratumoral treatment with STEAP1-BBζ CAR T cells was associated with significant tumor growth inhibition that was statistically significant by day 16 of treatment (**figure 3A**). Mice were sacrificed on day 25 and residual tumors from mice treated with STEAP1-BBζ CAR T cells showed large areas of necrotic debris and regions of viable tumor were infiltrated with CD3^+^ STEAP1-BBζ CAR T cells (**supplemental figure 6A**). STEAP1 expression was conserved in the tumors across treatment groups (**supplemental figure 6B**).

**Figure 3.**
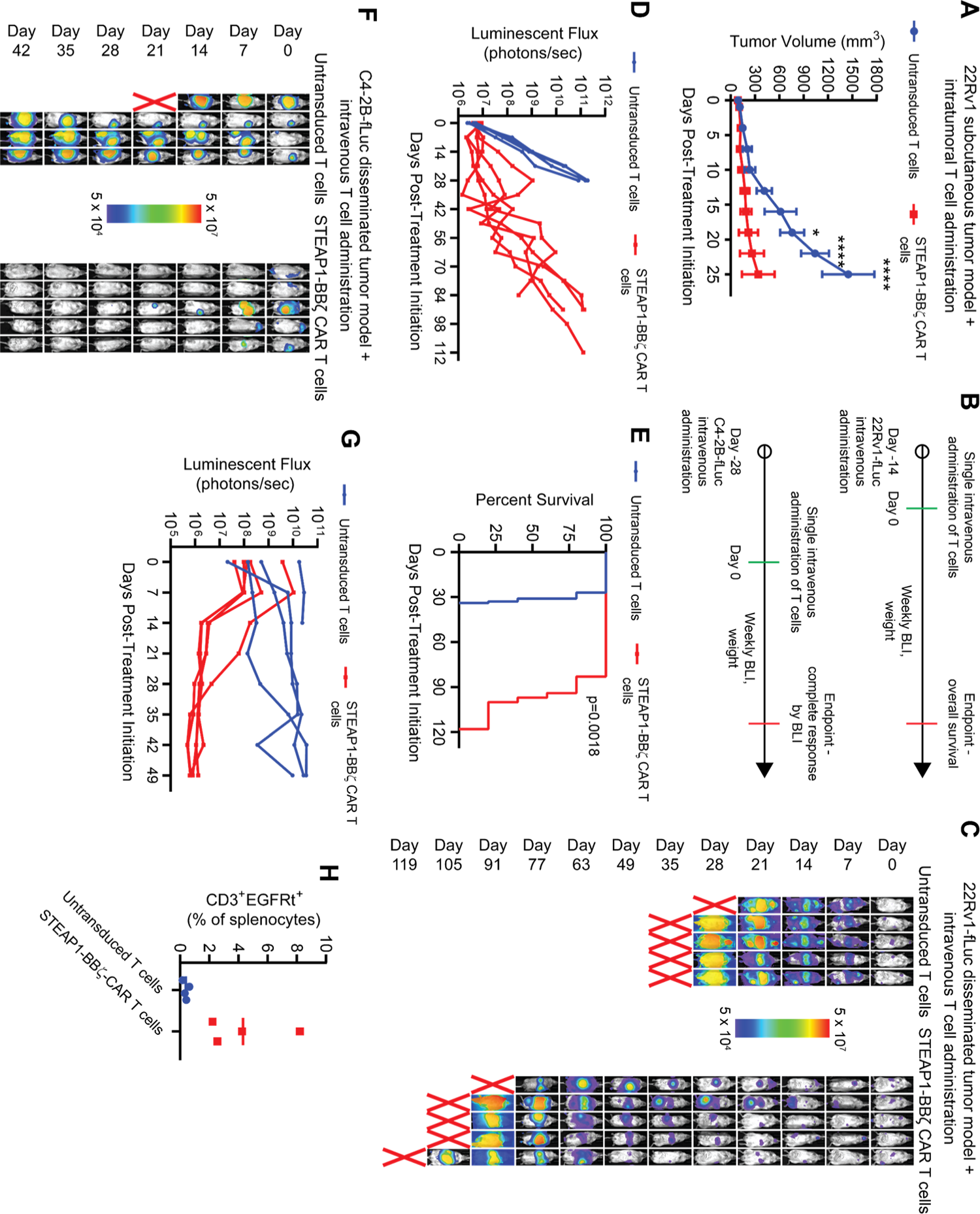
*In vivo* antitumor activity of STEAP1-BBζ CAR T cell therapy in prostate cancer models with native STEAP1 expression. (**A**) Volumes of 22Rv1 subcutaneous tumors in NSG mice over time after a single intratumoral injection of 5 x 10^6^ untransduced T cells or STEAP1-BBζ CAR T cells at normal CD4/CD8 ratios. (**B**) Schematic of tumor challenge experiments for 22Rv1 (top) and C4-2B (bottom) disseminated models. fLuc = firefly luciferase; BLI = bioluminescence imaging. (**C**) Serial live bioluminescence imaging (BLI) of NSG mice engrafted with 22Rv1-fLuc metastases and treated with a single intravenous injection of 5 x 10^6^ untransduced T cells or STEAP1-BBζ CAR T cells at normal CD4/CD8 ratios on day 0. Red X denotes deceased mice. Radiance scale is shown. (**D**) Plot showing the quantification of total flux over time from live BLI of each mouse in **C**. (**E**) Kaplan-Meier survival curves of mice in **C** with statistical significance determined by log-rank (Mantel-Cox) test. (**F**) Serial live BLI of NSG mice engrafted with C4-2B metastases and treated with a single intravenous injection of 5 x 10^6^ untransduced T cells or STEAP1-BBζ CAR T cells at normal CD4/CD8 ratios on day 0. Red X denotes deceased mice. Radiance scale is shown. (**G**) Plot showing the quantification of total flux over time from live BLI of each mouse in **F**. (**H**) Quantification of CD3^+^EGFRt^+^ STEAP1-BBζ CAR T cells by flow cytometry from splenocytes of mice treated with STEAP1-BBζ CAR T cells at the end of experiment on day 49. Error bars represent SD. * denotes p < 0.05; *** denotes p < 0.0001. For panel **A**, two-way ANOVA with Sidak’s multiple comparison test was used.

We transduced 22Rv1 cells with lentivirus to enforce firefly luciferase (fLuc) expression and 10^6^ 22Rv1-fLuc cells were injected into the tail veins of male NSG mice. Metastatic colonization was visualized by live bioluminescence imaging (BLI) after two weeks, at which point mice were treated with a single intravenous injection of either 5 x 10^6^ untransduced T cells or STEAP1-BBζ CAR T cells (**figure 3B**). Serial BLI revealed rapid disease progression in mice treated with untransduced T cells while those receiving STEAP1-BBζ CAR T cells demonstrated a significant delay in tumor progression (**figure 3C,D**) and extension of survival (97 days versus 31 days, p=0.0018 by log-rank test, **figure 3E**). There was no significant difference in mouse weights between treatment arms (**supplemental figure 6C**). IHC staining of tumors at the end of study showed a significant reduction in STEAP1 expression (**supplemental figure 6D,E**), indicating that antigen escape was a mechanism of resistance. However, this was unlikely a result of transdifferentiation to a variant prostate cancer state as we did not appreciate morphologic changes or loss of PSMA expression^49^ (**supplemental figure 6F**).

We also inoculated male NSG mice with C4-2B-fLuc cells by tail vein injection. C4-2B is a castration-resistant subline of LNCaP^50^ with growth kinetics more in line with typical prostate cancer. Four weeks after injection, metastatic colonization was confirmed by BLI and mice were treated with single intravenous injection of either 5 x 10^6^ untransduced T cells or STEAP1-BBζ CAR T cells (**figure 3B**). Serial BLI showed a complete response in all mice who received STEAP1-BBζ CAR T cells within five weeks of treatment (**figure 3F,G**). We identified a trend of increased weight loss in the untransduced T cell treatment group (**supplemental figure 7A**) but this was not statistically significant. Necropsy of mice treated with STEAP1-BBζ CAR T cells showed no macroscopic disease and *ex vivo* BLI of organs did not reveal any signal (**supplemental figure 7B**), suggesting that these mice were likely cured. We identified peripheral persistence of STEAP1-BBζ CAR T cells at the end of the experiment based on the presence of detectable CD3^+^EGFRt^+^ splenocytes (**figure 3H**).

### Mouse-in-mouse STEAP1 CAR T cell studies demonstrate antitumor therapeutic efficacy

The activation and cytolytic activity of STEAP1-BBζ CAR T cells observed in the very low STEAP1 antigen density (∼1,500 molecules/cell) context of the PC3 cell line (**figure 2G-I**, **supplemental figure 8A,B**) and evidence of *in vivo* antitumor activity in a disseminated PC3-fLuc tumor model (**supplemental figure 8C-E**) presented concerns about the potential for on-target, off-tumor toxicities. To evaluate for potential toxicity in a tractable model organism, we generated a human STEAP1 knock-in (hSTEAP1-KI) mouse in which the human *STEAP1* gene was knocked into the mouse *Steap1* gene locus on the C57Bl/6 background (**figure 4A**). A mouse colony was established with genotyping performed by polymerase chain reaction (PCR) of tail DNA (**figure 4B**). Both homozygous and heterozygous hSTEAP1-KI mice exhibited no apparent phenotypic or reproductive abnormalities compared to wildtype littermates. A tissue survey for human STEAP1 expression based on quantitative reverse transcription PCR (qRT-PCR) was performed on male and female heterozygous hSTEAP1-KI (hSTEAP1-KI/+) mice and revealed greatest relative expression in the prostate, followed by the uterus and adrenal gland (**figure 4C**). Further *in situ* analysis by STEAP1 IHC of male hSTEAP1-KI/+ prostate and adrenal glands revealed human STEAP1 expression confined to luminal epithelial cells of the prostate (**figure 4D**) and expression in the adrenal cortex (**figure 4E**).

**Figure 4.**
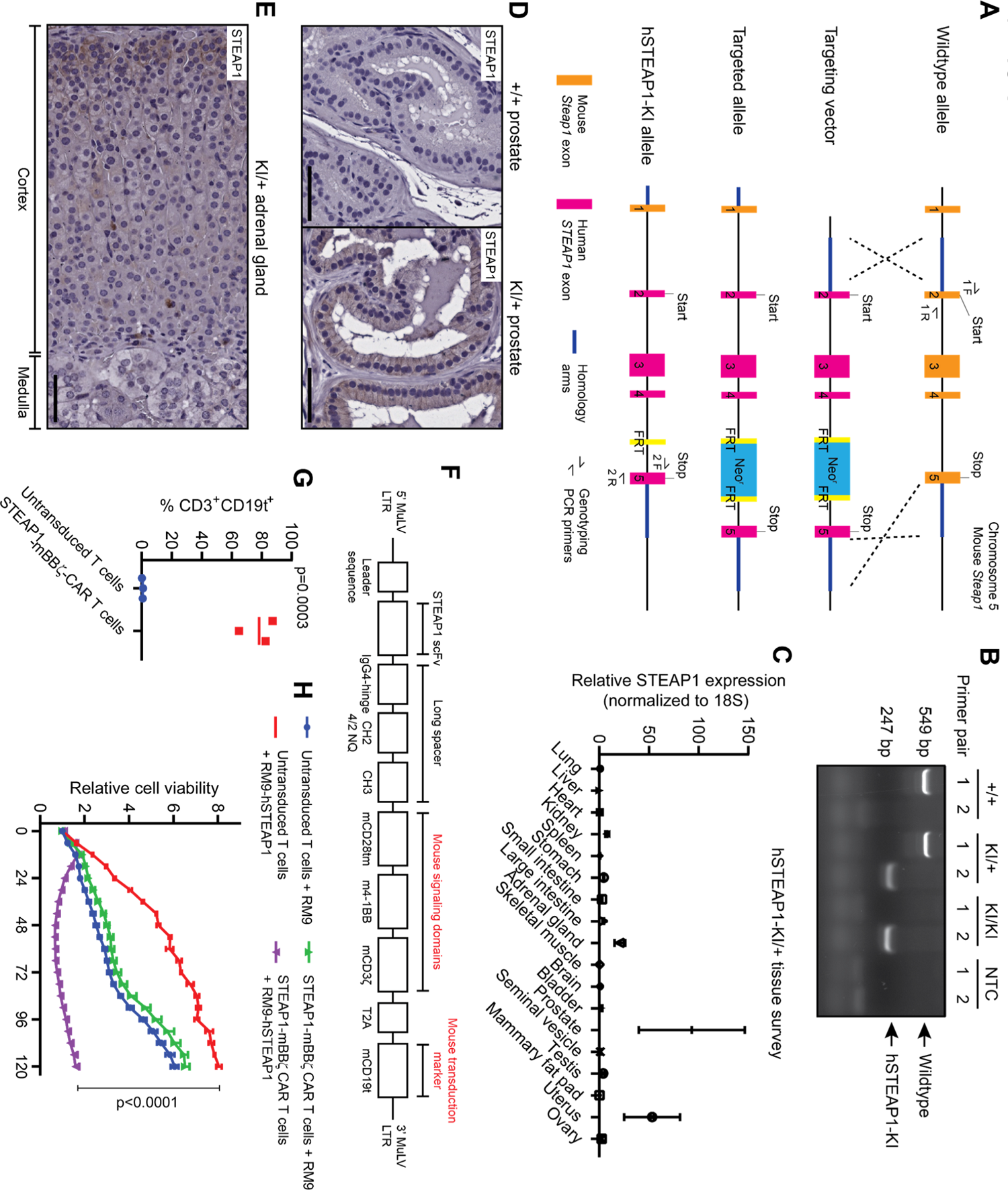
Establishing a mouse-in-mouse system with a novel human STEAP1 knock-in (hSTEAP1-KI) mouse model and murinized STEAP1 CAR. (**A**) Schematic showing the homologous recombination strategy using a targeting vector to knock-in human *STEAP1* exons 2-5 into the mouse *Steap1* locus on the C57Bl/6 background. FRT = Flippase recognition target. (**B**) Visualization of PCR products from tail tip genotyping of wildtype (+/+), heterozygous (KI/+), or homozygous (KI/KI) mice using primer pairs intended to amplify portions of wildtype or hSTEAP1-KI alleles. NTC = null template control. (**C**) qPCR for human STEAP1 expression normalized to 18S expression in a survey of tissues from hSTEAP1-KI/+ mice. n = 3 for sex-specific organs and n = 6 for common organs. Bars represent SD. Photomicrographs of STEAP1 IHC staining of (**D**) prostate tissues from (left) +/+ and (right) KI/+ mice and (**E**) an adrenal gland from a KI/+ mouse. Scale bars = 50 µm. (**F**) Schematic of the retroviral murinized STEAP1 CAR construct. MuLV = murine leukemia virus; mCD19t = mouse truncated CD19. (**G**) Quantification of the efficiency of retroviral transduction of activated mouse T cells from three independent experiments based the frequency of mouse CD3^+^CD19t^+^ cells by flow cytometry. (**H**) Relative cell viability of RM9 or RM9-hSTEAP1 target cells over time measured by fluorescence live cell imaging upon co-culture at a 1:1 ratio with mouse STEAP1-mBBζ CAR T cells or untransduced T cells. n = 4 replicates per condition. Error bars represent SEM. For panel **G**, unpaired two-tailed Student’s t test with Welch’s correction was used. In panel **H**, two-way ANOVA with Sidak’s multiple comparison test was used.

A murinized version of the STEAP1 CAR, called STEAP1-mBBζ CAR, in which the scFv and IgG4 hinge-CH2-CH3 spacer were retained but the CD28 transmembrane domain, 4-1BB costimulatory domain, and CD3ζ activation domain were replaced with their mouse orthologs was cloned into a gammaretroviral construct (**figure 4F**). In addition, the human EGFRt transduction marker was replaced with a truncated mouse CD19 (mCD19t) to minimize potential immunogenicity. We confirmed the efficient retroviral transduction of T cells enriched from mouse splenocytes (**figure 4G**) and demonstrated the capacity of mouse STEAP1-mBBζ CAR T cells to induce cytolysis of the RM9 mouse prostate cancer cell line^51^ engineered to express human STEAP1 (RM9-hSTEAP1) by lentiviral transduction (**figure 4H**).

The *in vivo* efficacy of mouse STEAP1-mBBζ CAR T cells was validated in a disseminated RM9-STEAP1-fLuc tumor model in NSG mice (**supplemental figure 9A**). One week after tail vein injection of RM9-STEAP1-fLuc cells, mice were treated with either 5 x 10^6^ untransduced mouse T cells or mouse STEAP1-mBBζ CAR T cells by tail vein injection. Mice that received untransduced mouse T cells demonstrated unchecked disease progression, whereas those treated with STEAP1-mBBζ CAR T cells uniformly exhibited rapid disease regression which was followed by subsequent relapse ten days later (**supplemental figure 9B,C**). STEAP1-mBBζ CAR T cell therapy was associated with a statistically significant survival benefit (22 days versus 12 days, p=0.0039 by log-rank test, **supplemental figure 9D**). Weight loss was evident in both treatment groups as tumor burden increased prior to death (**supplemental figure 9E,F**).

Analysis of mouse splenocytes collected at necropsy showed peripheral persistence of STEAP1-mBBζ CAR T cells with the detection of mCD3^+^ mCD19t^+^ cells up to 24 days after adoptive transfer (**supplemental figure 9G**). Lungs were harvested from mice in both treatment groups and STEAP1 IHC showed loss of STEAP1 expression in pulmonary metastases from mice treated with STEAP1-mBBζ CAR T cells (**supplemental figure 9H**).

We subsequently expanded clonal RM9-STEAP1-fLuc lines to determine whether the observed tumor antigen escape could be a result of pre-existing heterogeneity in STEAP1 expression. The experiment was repeated with a clonal, disseminated RM9-STEAP1-fLuc tumor model in NSG mice (**supplemental figure 10A**). In this context, mice treated with STEAP1-mBBζ CAR T cells demonstrated a prompt and durable complete response (**supplemental figure 10B-D**). These findings further highlight the potency of STEAP1-mBBζ CAR T cells in eradicating STEAP1^+^ prostate cancer and suggest that adjunct therapeutic strategies may be needed to overcome resistance in subgroups of advanced prostate cancer patients where inter- or intra-tumor STEAP1 heterogeneity is present (**figure 2B**).

### STEAP1 CAR T cell therapy is safe in a humanized STEAP1 mouse model

To investigate both the preclinical safety and efficacy of STEAP1-mBBζ CAR T cell therapy, we inoculated male heterozygous hSTEAP1-KI mice with syngeneic, non-clonal RM9-STEAP1-fLuc cells by tail vein injection (**figure 5A**). After confirmation of metastatic colonization by BLI about a week later, mice received pre-conditioning cyclophosphamide 100 mg/kg by intraperitoneal injection^52^. A day later, mice were randomized to treatment with either 5 x 10^6^ untransduced mouse T cells or mouse STEAP1-mBBζ CAR T cells by tail vein injection. All mice that received mouse STEAP1-mBBζ CAR T cells demonstrated a decrease in tumor burden within the first week of treatment initiation based on BLI (**figure 5B,C**). The observed response was short-lived but led to a modest extension of survival (21 days versus 12 days, p=0.0138 by log-rank test, **figure 5D**)—similar to findings from the non-clonal RM9-STEAP1-fLuc experiments in NSG mice (**supplemental figure 9D**).

**Figure 5.**
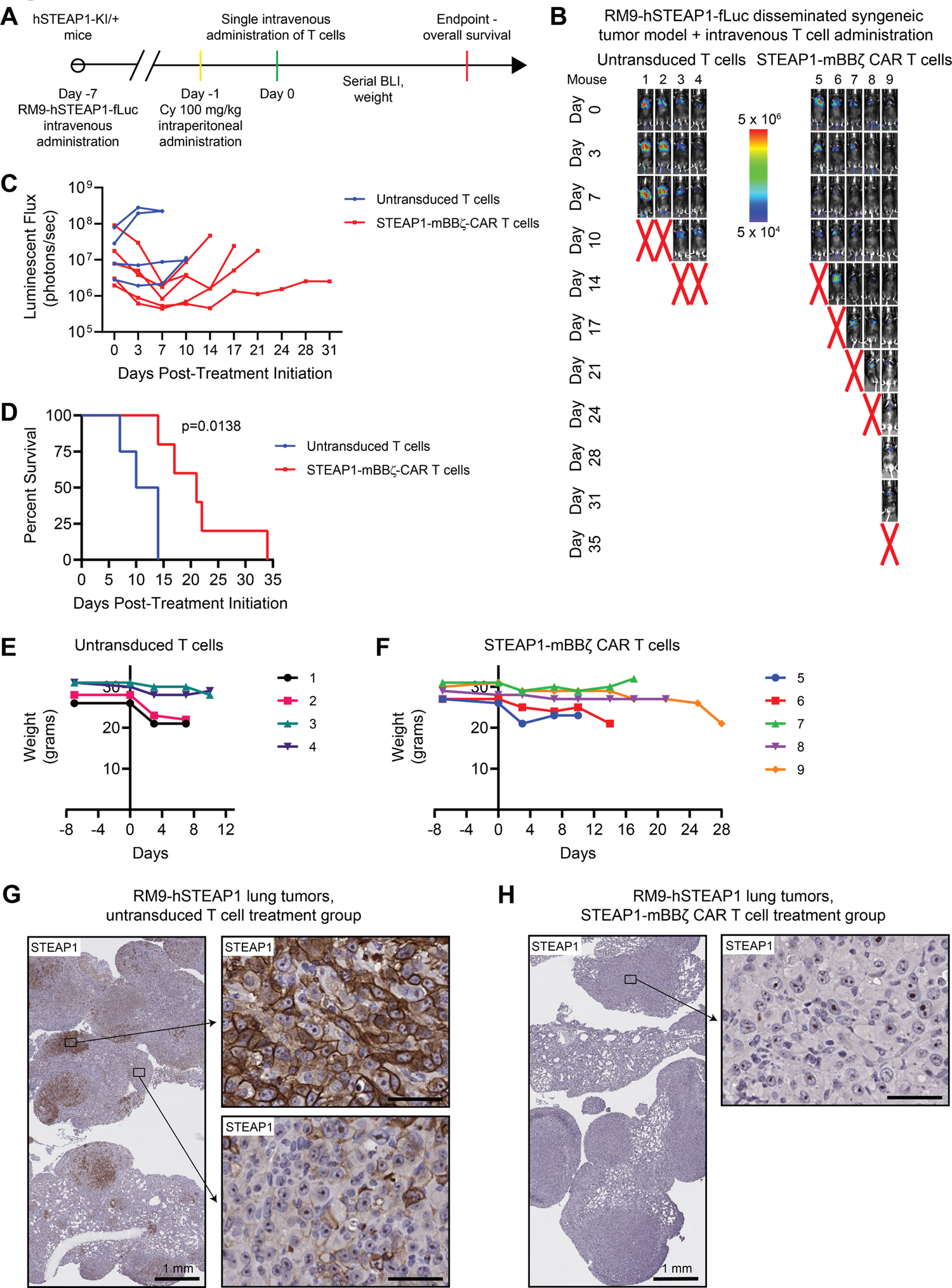
Determination of the efficacy and safety of mouse STEAP1-mBBζ CAR T cells in hSTEAP1-KI mice bearing syngeneic, disseminated prostate cancer. (**A**) Schematic of the tumor challenge experiment for the RM9-hSTEAP1 disseminated model in hSTEAP1-KI/+ mice. Cy = cyclophosphamide (for preconditioning). (**B**) Serial live BLI of hSTEAP1-KI/+ mice engrafted with RM9-hSTEAP1-fLuc metastases and treated with a single intravenous injection of 5 x 10^6^ mouse untransduced T cells or STEAP1-mBBζ CAR T cells on day 0. Red X denotes deceased mice. Radiance scale is shown. (**C**) Plot showing the quantification of total flux over time from live BLI of each mouse in **B**. (**D**) Kaplan-Meier survival curves of mice in **B** with statistical significance determined by log-rank (Mantel-Cox) test. Plots of weights for each mouse (numbered in **B**) over time in the (**E**) mouse untransduced T cell treatment group and (**F**) STEAP1-mBBζ CAR T cell treatment group. (**G**) Photomicrographs at (left) low and (right) high magnification of STEAP1 IHC staining of RM9-hSTEAP1 lung tumors after treatment with mouse untransduced T cells showing regions of strong homogenous STEAP1 expression and heterogeneous STEAP1 expression. Scale bars = 50 µm, unless otherwise noted. (**H**) Photomicrographs at (left) low and (right) high magnification of STEAP1 IHC staining of RM9-hSTEAP1 tumors after treatment with STEAP1-mBBζ CAR T cells showing no STEAP1 expression. Scale bars = 50 µm, unless otherwise noted.

There were no gross toxicities or premature deaths specifically associated with mouse STEAP1-mBBζ CAR T cell therapy at this dose level where clear evidence of antitumor efficacy was observed. Weight loss was associated with increased tumor burden but common to both treatment arms (**figure 5E,F**). Plasma cytokine analysis of IFN-γ, IL-2, IL-6, and TNF-α at day 0 and day 8 of treatment showed no changes indicative of cytokine storm (**supplemental figure 11**). Importantly, heterozygous hSTEAP1-KI mice treated with STEAP1-mBBζ CAR T cells demonstrated no obvious tissue disruption or increased infiltration of CD3^+^ T cells in the prostate (**supplemental figure 12A,B**) or adrenal gland (**supplemental figure12C,D**) relative to their counterparts treated with untransduced T cells. Lungs collected at the end of the experiment showed human STEAP1 expression in pulmonary metastases with regional heterogeneity in mice treated with untransduced mouse T cells (**figure 5G**). On the other hand, tumors from mice treated with mouse STEAP1-mBBζ CAR T cells again demonstrated an absence of human STEAP1 expression (**figure 5H**). These data provide preliminary preclinical evidence that STEAP1 can be safely targeted with potent CAR T cell therapy without severe toxicities.

## Discussion

The effectiveness of CAR T cell therapy and other immune-based targeted therapeutics is highly dependent on consistent antigen expression on all or most cells comprising the tumor population within an individual patient. However, antigen heterogeneity is pronounced in solid tumors including prostate cancer, where progression to mCRPC and treatment resistance are associated with the emergence of divergent disease subtypes marked by distinct transcriptional programs^53–55^ and cell surface antigen expression^13^. While PSMA is considered one of the foremost biomarkers in prostate cancer with significant overexpression found across the spectrum of disease progression, our work corroborates findings from a recent publication^10^ indicating that PSMA expression is heterogeneous in lethal mCRPC. We show that STEAP1 is more broadly expressed than PSMA in this setting but is by no means expressed uniformly at high levels in all mCRPC tissues. No single antigen-targeted therapy including CAR T cell therapy may be able to overcome pre-existing tumor antigen heterogeneity in mCRPC. Thus, it is of critical importance to thoroughly credential additional therapeutic targets such as STEAP1 in mCRPC that may enable combinatorial therapies that exert insurmountable therapeutic pressure. These include dual antigen-targeted (e.g., PSMA and STEAP1) CAR T cell therapies or multimodal strategies combining CAR T cell therapies with ADCs, T-BsAbs, or other treatments that potently promote antigen-independent and -dependent tumor killing.

We engineered a STEAP1-targeted CAR T cell therapy that is highly antigen-specific and functionally localized the epitope recognized by the CAR to the second ECD of STEAP1. Our STEAP1 CAR T cells demonstrate substantial antitumor activity against multiple disseminated prostate cancer models both in human-in-mouse and mouse-in-mouse studies. Importantly, our STEAP1 CAR is capable of inducing T cell activation and target cell cytolysis even in low antigen density conditions, as evidenced by reactivity against the PC3 prostate cancer model.

However, this sensitivity of STEAP1 CAR T cells to low levels of STEAP1 expression may be advantageous from the perspective of enhancing antitumor efficacy but could also accentuate liabilities from on-target off-tumor toxicity. Systemic expression of STEAP1 has previously been reported as virtually absent in normal human tissues^14, 56^ except the prostate gland where membranous expression in prostate epithelial cells has been described^24^. To delve the safety of STEAP1 CAR T cell therapy in the preclinical setting, we exceeded the standard in the field by generating a humanized STEAP1 mouse model. The hSTEAP1-KI mouse model recapitulated human STEAP1 expression in the prostate gland and showed expression in the adrenal cortex. Reassuringly, STEAP1 CAR T cell therapy at a dose sufficient to induce antitumor activity did not lead to evident systemic toxicities in hSTEAP1-KI mice including on-target off-tumor toxicities at sites of human STEAP1 expression.

A recurring mechanism of prostate cancer relapse and progression after STEAP1 CAR T cell therapy in our studies was tumor antigen escape. On one hand, this finding underscores the overall potency of our STEAP1 CAR T cell therapy. However, it is unclear whether the loss of tumor STEAP1 expression is solely due to inherent tumor antigen heterogeneity or whether there is also adaptive downregulation of STEAP1 expression. A recent publication showed that promoter methylation of *STEAP1* modulates STEAP1 expression and epigenetic deregulation by DNA methyltransferase and histone deacetylase inhibition was sufficient to significantly upregulate STEAP1 expression^57^. Perhaps treatment with epigenetic inhibitors in combination with STEAP1 CAR T cell therapy could simultaneously enhance tumor STEAP1 expression and reprogram CAR T cells to favorable exhaustion-resistant differentiation states^58, 59^, thereby mitigating tumor antigen loss and enhancing antitumor efficacy in prostate cancer. Another consideration is that STEAP1 has been functionally implicated in cancer progression and a prior study indicated that acute STEAP1 gene knockdown reduced cell viability and proliferation while inducing apoptosis in prostate cancer^34^. However, the mechanistic basis for these effects has not been elucidated. An interesting observation is that STEAP1 is unique from other STEAP family members (STEAP2, 3, and 4) in that it lacks an intracellular oxidoreductase domain^18^ which is necessary for metalloreductase activity. As a result, STEAP1 homotrimers, but not heterotrimers with other STEAP proteins, lack enzymatic function to reduce Fe3+ to Fe2+ and Cu2+ to Cu1+. Whether and how the involvement of STEAP1 in metal ion metabolism promotes cancer progression has yet to be determined and is worthy of further investigation.

The immunologically ‘cold’ tumor microenvironment of prostate cancer is a major barrier to the efficacy of cancer immunotherapies including immune checkpoint inhibitors. In addition, exploratory studies associated with a phase I clinical trial of PSMA CAR T cell therapy armored to express dominant-negative transforming growth factor-β receptor (TGFβR-DN) in mCRPC showed that the expression of immunosuppressive signaling molecules in the tumor microenvironment increases after CAR T infusion^9^. In our work, we have used a disseminated, syngeneic RM9-hSTEAP1 tumor model in hSTEAP1-KI to approximate the immunosuppressive nature of mCRPC based on prior characterization of RM9 as a poorly immunogenic model^52, 60^. One caveat is that this may not fully recapitulate the complexity of the human tumor immune microenvironment. Thus, additional approaches such as armoring STEAP1 CAR T cells to express TGFβR-DN^61^ or recombinant cytokines^62^ (e.g., IL-12, IL-15, or IL-18) or the concurrent systemic administration of novel immunomodulators may be necessary to enhance the effector function of STEAP1 CAR T cells within the hostile tumor microenvironment of prostate cancer.

The findings of our studies have led to a partnership with the National Cancer Institute (NCI) Experimental Therapeutics (NExT) Program with the goal of translating STEAP1 CAR T cell therapy to a first-in-human trial for men with mCRPC. Safety and efficacy signals from this early phase clinical trial will help determine whether there may also be value in investigating this therapeutic approach for other cancer types that highly express STEAP1.

## Methods

### Cell lines

22Rv1 (CRL-2505), LNCaP (CRL-1740), PC3 (CRL-1435), DU145 (HTB-81), NCI-H660 (CRL-5813), C4-2B (CRL-3315), RM9 (RL-3312), and Myc-CaP (CRL-3255) were obtained from the American Type Culture Collection. LNCaP95 cells were a gift from Stephen R. Plymate (University of Washington, Seattle). MSKCC EF1 were derived from the MSKCC PCa4 organoid line provided by Yu Chen (Memorial Sloan Kettering Cancer Center), as previously described^13^. Cell lines were maintained in RPMI 1640 medium supplemented with 10% FBS, 100 U/mL penicillin and 100 µg/mL streptomycin, and 4 mmol/L GlutaMAX (Thermo Fisher). 22Rv1 STEAP1 ko and PC3 STEAP1 ko cells were generated by transient transfection of 22Rv1 cells with a pool of PX458 (Addgene, #48138) plasmids each expressing one of four different sgRNA targeting sequences predicted from the Broad Institute Genetic Perturbation Platform sgRNA Designer^63^: 1) 5’-ATAGTCTGTCTTACCCAATG-3’; 2) 5’-CCTTTGTAGCATAAGGACAC-3’; 3) 5’-ATCCACTTATCCAACCAATG-3’; and 4) 5’-CATCAACAAAGTCTTGCCAA-3’. 48-72 hours after transfection, GFP-positive cells were singly sorted on a Sony SH800 Cell Sorter into a 96-well plate and clonally expanded.

gBlocks encoding firefly luciferase (fLuc), hSTEAP1, mSTEAP1, mSTEAP1 hECD1, mSTEAP1 hECD2, mSTEAP1 hECD3, hSTEAP1B isoform 1, hSTEAP1B isoform 2, and hSTEAP1B isoform 3 were cloned into the EcoRI site of FU-CGW^64^ by HiFi DNA Assembly. The FU-hSTEAP1-CGW plasmid was modified to excise the GFP cassette by digestion with AgeI and BsrGI. A PCR product encoding firefly (fLuc) was then inserted by HiFi DNA assembly to generate the FU-hSTEAP1-C-fLuc-GW plasmid. These lentiviruses were produced and titered as previously described^65^. Lentiviruses were used to introduce stable transgene expression in the cell lines noted above.

### Animal studies

All mouse studies were performed in accordance with protocols approved by the Fred Hutchinson Cancer Center Institutional Animal Care and Use Committee and regulations of Comparative Medicine.

For studies using immunocompromised mice, six- to eight-week-old male NSG (NOD-SCID-IL2Rγ-null) mice were obtained from The Jackson Laboratory. For the 22Rv1 subcutaneous tumor and intratumoral T cell administration model, 2 x 10^6^ 22Rv1 cells were suspended in 100 µl ice-cold Matrigel matrix (Corning) and injected subcutaneously into the flanks of NSG mice. Tumors were measured twice weekly using electronic calipers and tumor volume (TV) calculated based on the equation TV = ½ (L * W^2^). When TV was ∼75-100 mm^3^, 5 x 10^6^ untransduced or STEAP1 CAR T cells at a defined CD4/CD8 composition of 1:1 and suspended in 100 µl of PBS was injected intratumorally. Mice were sacrificed 25 days after intratumoral T cell therapy. For disseminated human prostate cancer and intravenous T cell administration models, between 5 x 10^5^ and 10^6^ prostate cancer cells were suspended in 100 µl of PBS and injected into the tail veins of NSG mice. Tumor burden was monitored by live bioluminescence imaging on an IVIS Spectrum (PerkinElmer) after intraperitoneal injection of XenoLight D-luciferin (PerkinElmer). When metastatic colonization was confirmed by imaging, 5 x 10^6^ human untransduced or STEAP1-BBζ CAR T cells at a defined CD4/CD8 composition of 1:1 and suspended in 100 µl of PBS was injected by tail vein. Metastatic tumors and spleen were harvested when mice were euthanized at compassionate endpoints. In the disseminated C4-2B model, lungs and livers were collected from a subset of mice and placed individually in six-well plates in DMEM media supplemented with 10% FBS and GlutaMAX. *Ex vivo* bioluminescence imaging was performed by introducing XenoLight D-luciferin into the media and quantifying signal on an IVIS Spectrum. Metastatic tumors were formalin-fixed and paraffin-embedded. Splenocytes were harvested from spleens to perform flow cytometry to evaluate the peripheral persistence of CAR T cells.

For animal studies using hSTEAP1-KI mice, heterozygous hSTEAP1-KI mice were generated by crossing homozygous hSTEAP1-KI mice with wildtype C57Bl/6 mice. Genotyping of all hSTEAP1-KI mice was performed by PCR of 10 ng of tail DNA using the Taq 2X Master Mix (New England Biolabs) and visualization of PCR products by gel electrophoresis on a 2% agarose gel. Primers used for genotyping PCR reactions are 1) wildtype: forward 5’-CTAGGTGGCTGAAGCCGTA-3’ and reverse 5’-GCGATGACCAAAAGTGACTTC-3’, 2) hSTEAP1-KI: forward 5’-CAGATGAGGTAGGATGGGATAAAC-3’ and reverse 5’-CCTCAAGCATGGCAGGAATAG-3’. Thermocycler conditions for genotyping PCR are 95°C x 30 seconds; (95°C x 30 seconds, 58°C x 30 seconds, 68°C x 70 seconds) x 35 cycles; 68°C x 5 minutes; and 12°C hold.

For disseminated RM9-hSTEAP1-fLuc mouse prostate cancer and intravenous T cell administration models, 5 x 10^5^ RM9-hSTEAP1-fLuc cells were suspended in 100 µl of PBS and injected into the tail veins of either NSG or heterozygous hSTEAP1-KI mice. Tumor burden was monitored by live bioluminescence imaging on an IVIS Spectrum (PerkinElmer) after intraperitoneal injection of XenoLight D-luciferin (PerkinElmer). When metastatic colonization was confirmed by imaging, 5 x 10^6^ mouse untransduced or STEAP1-mBBζ CAR T cells suspended in 100 µl of PBS was injected by tail vein. Retroorbital blood was collected using heparinized capillary tubes into polystyrene tubes containing an EDTA/PBS solution. After collection, retroorbital blood samples were incubated at room temperature for 15-20 minutes and centrifuged in a tabletop centrifuge at 2,000 x g for 10 minutes. Plasma was collected and stored at −80°C. Lungs, spleen, prostate, and adrenal glands were harvested when mice were euthanized at compassionate endpoints. Lungs, prostate, and adrenal glands were formalin-fixed and paraffin-embedded. Splenocytes were harvested from spleens to perform flow cytometry to evaluate the peripheral persistence of CAR T cells.

### Immunohistochemical studies

Tissue sections were deparaffinized in xylene and rehydrated in 100%, 95%, 75% ethanol, and finally TBS with 0.1% Tween 20 (TBST). Antigen retrieval was conducted in Citrate-Based Antigen Unmasking Solution (Vector Labs) using a pressure cooker at 95°C for 30 minutes. Tissue sections were blocked with Dual Endogenous Enzyme-Blocking Reagent (Agilent Technologies) and incubated for 10 minutes followed by three washes with TBST. Slides were incubated with primary antibody in a humidified chamber at 37°C for one hour. Primary antibodies and dilutions used for this study are rabbit anti-STEAP1 antibody (LS Bio, LS-C291740, 1:500), rabbit anti-CD3 antibody (Thermo Fisher, MA5-14524, 1:100), and mouse anti-PSMA antibody (Dako, M3620, 1:50). Slides were washed with TBST and incubated with PowerVision Poly-HRP anti-rabbit IgG or anti-mouse IgG (Leica Biosystems) in a humidified chamber at 37°C for 30 minutes. Slides were washed with TBST and incubated with 3,3′-Diaminobenzidine (DAB) (Sigma Aldrich) for at room temperature for 10 minutes. DAB was quenched in deionized water. Slides were stained in Dako hematoxylin (Agilent Technologies) at room temperature for one minute and washed with deionized water for five minutes. Slides were dehydrated in 75%, 95%, 100% ethanol, and finally xylene. Slides were mounted using Permount mounting medium (Fisher Chemical) and cover slipped.

### Prostate cancer tissue microarray analysis

University of Washington mCRPC Tissue Acquisition Necropsy (TAN) tissue microarrays (Prostate Cancer Biorepository Network) were used for immunohistochemical studies. Each core stained with either STEAP1 or PSMA was scored by an experienced pathologist (M.P.R) and assigned an intensity score of 0, 1, 2, or 3 and frequency of positive cell staining ranging from 0% to 100%. H-scores were generated for each core by multiplying the intensity score by the frequency of positive cell staining resulting in a minimum of 0 and maximum of 300. The average H-score of replicate cores represented in the tissue microarray was determined for each mCRPC tissue.

### Immunoblotting

Protein extracts were collected in 9 M urea lysis buffer and quantified using the Pierce Rapid Gold BCA Protein Assay Kit (Thermo Fisher). Protein samples were fractionated with SDS-PAGE using Bolt 4-12% Bis-Tris Plus Gels (Thermo Fisher) and transferred to nitrocellulose membranes using the Invitrogen Mini Blot module (Thermo Fisher). Membranes were blocked with 5% non-fat milk in PBS + 0.5% Tween 20 (PBST) on a shaker at room temperature for 30 minutes. Membranes were then incubated with primary antibody on a shaker at 4°C overnight. Primary antibodies used for this study are mouse anti-STEAP (Santa Cruz, sc-271872, 1:1,000) and GAPDH (GeneTex, GX627408, 1:5,000). Membranes were washed with PBST and incubated with goat anti-mouse IgG (H+L) secondary antibody conjugated with horseradish peroxidase (Thermo Fisher, 31430, 1:10,000) on a shaker at room temperature for 1 hour. Membranes were washed with PBST, incubated with Immobilon Western Chemiluminescent HRP Substrate (EMD Millipore) at room temperature for three minutes, and visualized on a ChemiDoc MP Imaging System (Bio-Rad Laboratories).

### Absolute STEAP1 quantification

Flow cytometric quantification of STEAP1 antigen density across human prostate cancer cell lines was performed using Quantum Simply Cellular microspheres (Bangs Laboratories) per the manufacturer’s protocol. Vandortuzumab (Creative Biolabs) was used as the primary antibody and rat anti-human IgG Fc antibody conjugated to APC (BioLegend) was used as the secondary antibody. Stained cells and beads were analyzed on a BD FACSCanto II (BD Biosciences). The Geo Mean or Median channel values for each population were recorded into the provided QuickCal spreadsheet yielding a regression coefficient of 0.998.

### Chimeric antigen receptors (CAR) expression plasmids

gBlocks (Integrated DNA Technologies) encoding the GM-CSF leader, DSTP3086S scFv (VL-[G4S]3-VH), IgG4 spacers, CD28 transmembrane domain, 4-1BB costimulatory domain, CD3ζ chain, EGFRt or mCD19t, and WPRE were cloned into the EcoRI site of pCCL-c-MNDU3-X (gift from Donald Kohn, Addgene plasmid #81071) or pMYs (Cell Biolabs) by HiFi DNA Assembly (New England Biolabs). Sequences involving cloning junctions and open reading frames were validated by Sanger sequencing at the Fred Hutch Genomics & Bioinformatics Shared Resource.

### CAR lentivirus and retrovirus production

For STEAP1-BBζ CAR lentivirus production, HEK 293T cells (ATCC) were thawed, cultured, and expanded in DMEM media supplemented with 10% FBS and GlutaMAX. HEK 293T cells were seeded on plates coated with Cultrex Poly-L-Lysine (R&D Systems) prior to transfection with the pCCl-c-MNDU3 STEAP1-BBζ CAR lentiviral plasmid and the packaging plasmids pMDL, pVSVg, and pREV using the TransIT-293 transfection reagent (Mirus Bio). About 18 hours after transfection, sodium butyrate and HEPES were added to each plate to a final concentration of 20 mM each. Eight hours later, media was aspirated from the plates and each plate was washed with PBS. DMEM media supplemented with 10% FBS, GlutaMAX, and 20 mM HEPES was added to each plate. Lentiviral supernatant was collected at 48 hours after transfection, vacuum filtered through a 0.22 µm filter, and concentrated by ultracentrifugation in polypropylene Konical tubes (Beckman Coulter) at 22,000 rpm at 4°C for two hours in an Optima XE 90 (Beckman Coulter). Lentiviral pellets were resuspended in the minimal residual media present after aspirating off supernatant, aliquoted in cryovials and stored at −80°C.

For STEAP1-mBBζ CAR retrovirus production, PLAT-E cells (Cell Biolabs) were thawed and cultured in DMEM media supplemented with 10% FBS, GlutaMAX, 1 µg/ml puromycin, and 10 µg/ml blasticidin. One day prior to seeding cells for transfection, PLAT-E cells were washed and seeded in antibiotic-free DMEM media supplemented with 10% FBS and GlutaMAX. PLAT-E cells were transfected with the pMYs STEAP1-mBBζ CAR retroviral construct using the FuGENE HD transfection reagent (Promega). 48 and 72 hours after transfection, supernatants containing retrovirus were passed through a 0.22 µm syringe filter prior to use in transduction.

### Human CAR T cell manufacturing

Peripheral blood mononuclear cells (PBMCs) from three de-identified healthy donors obtained by pheresis from the Fred Hutch Co-Operative Center for Excellence in Hematology were thawed and washed with pre-warmed TCM base media consisting of AIM-V media (Gibco) supplemented with 55 mM beta-mercaptoethanol, human male AB plasma (Sigma), and GlutaMAX. PBMCs were centrifuged in a tabletop centrifuge at 1500 rpm for 5 minutes. Cells were resuspended in TCM base media and counted on a hemacytometer. Dynabeads CD8 and CD4 Positive Isolation Kits were used per manufacturer’s protocol to separate CD8 and CD4 T cells. After bead detachment, CD8 T cells were seeded in CD8 media (TCM base media supplemented with 50 U/ml human IL-2 and 0.5 ng/ml human IL-15) and CD4 T cells were seeded in CD4 media (TCM base media supplemented with 0.5 ng/ml human IL-15 and 5 ng/ml human IL-7). CD8 and CD4 T cells were activated and expanded with Dynabeads Human T-Activator CD3/CD28 (Thermo Fisher) per manufacturer’s protocol. After two to four days, CD8 and CD4 T cells were counted and transduced with STEAP1-BBζ CAR lentivirus at a relative multiplicity-of-infection of 10 based on the infectious titer on HEK 293T cells. Lentiviral transduction was performed in the presence of 10 µg/ml protamine sulfate. 48 hours after transduction, T cells were collected, activation beads removed, and transduction efficiency of the T cells evaluated by flow cytometry. CAR-modified CD4 and CD8 T cells were counted every two days and maintained at a density of 10^6^ cells/ml in their respective CD4 and CD8 media.

### Mouse CAR T cell manufacturing

Splenocytes were harvested from the manual dissociation of spleens obtained from heterozygous hSTEAP1-KI mice. Splenocytes were passed through a 70 µm strainer and pelleted by centrifugation at 1,600 rpm for six minutes. Cells were resuspended in RBC lysis buffer (BioLegend) and incubated on ice for five minutes. Cells were washed with PBS and pelleted by centrifugation. Murine CD3 T cells were isolated using Mouse CD3+ T Cell Enrichment Columns (R&D Systems) per the manufacturer’s protocol. T cells were cultured in RPMI 1640 media containing 10% FBS, 50 U/ml human IL-2, 10 ng/ml murine IL-7, and 50 µM beta-mercaptoethanol. T cells were activated and expanded with Dynabeads Mouse T-Activator CD3/CD28 (Thermo Fisher) per manufacturer’s protocol. 48 and 72 hours later, mouse T cells were transduced with filtered pMYs STEAP1-mBBζ CAR retroviral supernatants via spinoculation on a tabletop centrifuge at 2,000 x g at 30°C for two hours. On day six of culture, beads were magnetically removed and T cell transduction efficiency was determined by flow cytometry prior to use in functional assays.

### Immunophenotyping CAR T cell products and assessment of peripheral persistence

1.5 x 10^5^ human PBMCs, human untransduced or STEAP1-BBζ CAR T cells, mouse untransduced or STEAP1-BBζ CAR T cells, or splenocytes were incubated with fluorophore conjugated antibodies on ice for 20 minutes. Antibodies used for human cells are mouse anti-human CD3 conjugated to APC (Thermo Fisher, 47-0036-42), mouse anti-human CD8 conjugated to FITC (BD Biosciences, 555366), rabbit anti-human EGFR (cetuximab) conjugated to PE (Novus Biologicals, NBP2-52671PE), mouse anti-human CD3 conjugated to BUV395 (BD Biosciences, 563548), mouse anti-human CD4 conjugated to BV605 (BioLegend, 344645), mouse anti-human CD8 conjugated to BUV805 (BD Biosciences, 564912), mouse anti-human CD45RO conjugated to BV510 (BioLegend, 304246), mouse anti-human CD45RA conjugated to BV711 (BioLegend, 304138), mouse anti-human PD-1 conjugated to BV421 (BD Biosciences, 565935), mouse anti-human CD95 conjugated to BUV615 (BD Biosciences, 752346), mouse anti-human CXCR3 conjugated to BV421 (BioLegend, 353716), mouse anti-human CD62L conjugated to BV785 (BD Biosciences, 565311), and mouse anti-human LAG-3 conjugated to BV421 (BD Biosciences, 565721). Antibodies used for mouse cells are rat anti-mouse CD8a conjugated to FITC (BioLegend, 100706) and rat anti-mouse CD19 conjugated to PE (BioLegend, 115508). Cells were washed with PBS after antibody staining and acquired on a BD FACSCanto II or BD Symphony 4. Data were analyzed on FlowJo v.10 (Treestar).

### Immunologic co-culture assays

CAR T cell functional assays were performed by co-culturing prostate cancer cells engineered to express GFP with either human or mouse untransduced or STEAP1 CAR T cells at variable effector-to-target (E:T) ratios in 96-well plates. For cytotoxicity assays, 96-well clear bottom black wall plates (Corning) were coated with Cultrex Poly-L-Lysine for 30 minutes and seeded with prostate cancer cells. Plates were incubated at 37°C for one hour. Effector cells were then counted and seeded into wells with tumor cells at specified E:T ratios. The plates were placed into a BioTek BioSpa 8 Automated Incubator (Agilent Technologies) and read by brightfield and fluorescence imaging on a BioTek Cytation 5 Cell Imaging Multi-Mode Reader (Agilent Technologies) every six hours for a total of six days. Target cells were quantified based on the number of GFP^+^ objects identified per scanned area using BioTek Gen5 Imager Software (Agilent Technologies). To assess T cell activation based on cytokine release, 25-50 µl of co-culture supernatants were collected at 24 and 48 hours of co-culture and stored at −30°C.

### Mitogen stimulation of T cells

T cells were stimulated with 5 ng/ml of phorbol 12-myristate 13-acetate (PMA, Sigma Aldrich) and 250 ng/ml of ionomycin (Sigma Aldrich) or with 5 µg/ml of phytohemagglutinin-L (PHA-L, Sigma Aldrich). Supernatants were collected at 24 hours for use as a positive control for IFN-γ ELISA studies or T cells were used as a positive control for induction of the exhaustion markers PD-1 and LAG-3 as assessed by flow cytometry.

### Enzyme-linked immunosorbent assay

To determine IFN-γ levels in co-culture supernatants, samples were thawed and sandwich ELISA for human or mouse IFN-γ levels was performed using the BD Human IFN-gamma ELISA Set (BD Biosciences, 555142) or BD Mouse IFN-gamma ELISA Set (BD Biosciences, 555138) according to the manufacturer’s protocol. Plates were read at 450 nm and 560 nm wavelengths using a BioTek Cytation 3 Cell Imaging Multi-Mode Microplate Reader (Agilent Technologies). Plasma samples isolated from retroorbital bleeds were used for ProcartaPlex immunoassays (Thermo Fisher) to quantify levels of mouse IFN-γ, IL-2, IL-6, and TNF-α according to the manufacturer’s protocol. Samples were assayed on a Luminex 100/200 System (Luminex).

## Data availability

All data related to the study are included in the article or uploaded as supplementary information.

## Author contributions

V.B.: Data curation, formal analysis, validation, investigation, visualization, methodology, writing-original draft, writing-review and editing. N.V.K.: Data curation, formal analysis, investigation, visualization, and methodology. T.E.P.: Conceptualization, data curation, formal analysis, investigation, visualization, and methodology. L.W.: Data curation and investigation. A.T.: Data curation and investigation. K.S.: Data curation and investigation. L.T.W.: Data curation and investigation. A.Z.: Data curation and investigation. D.R.: Data curation and investigation. R.G.: Formal analysis. R.P.: Data curation and investigation. M.P.R.: Formal analysis. L.T.: Data curation and writing-review and editing. M.C.H.: Data curation, formal analysis, and writing-review and editing. P.N.S.: Conceptualization and writing-review and editing. S.J.P.: Formal analysis, methodology, writing-review and editing. J.I.: Formal analysis, validation, investigation, and writing-review and editing. J.K.L.: Conceptualization, data curation, formal analysis, validation, investigation, visualization, methodology, writing-original draft, writing-review and editing.

## Acknowledgments

We grateful first and foremost to the patients and their families for their contributions, without which this research would not have been possible. We acknowledge Celestia Higano, Evan Yu, Heather Chang, Bruce Montgomery, Elahe Mostaghel, Andrew Hsieh, Daniel Lin, Funda Vakar-Lopez, Xiaotun Zhang, Lawrence True, and the rapid autopsy teams for the contributions to the University of Washington Prostate Cancer Donor Rapid Autopsy Program. We thank the Fred Hutch Co-Operative Center for Excellence in Hematology (supported by U54 DK106829). We also thank the Fred Hutch Genomics Shared Resource, Comparative Medicine Shared Resource, Flow Cytometry Shared Resource, Experimental Histopathology Shared Resource, and Immune Monitoring Shared Resource (supported by NIH/NCI Cancer Center Support Grant P30 CA015704). This research was funded in part by a Department of Defense Prostate Cancer Research Program Award (W81XWH-21-1-0581 to J.K.L.), a Fred Hutch/University of Washington Cancer Consortium Safeway Pilot Award (J.K.L.), a Seattle Cancer Care Alliance/Swim Across America Award (J.K.L.), the Pacific Northwest Prostate Cancer SPORE P50 CA097186 (P.S.N., L.T., and R.G.), the Institute for Prostate Cancer Research (M.P.R), the Doris Duke Charitable Foundation (2021184 to M.C.H), NCI R50 CA221836 (R.G.), JSPS Overseas Research Fellowships (202160429 to K.S.) and a Movember Foundation-Prostate Cancer Foundation Challenge Award (P.S.N. and J.K.L.).

