## Supplemental Data for "Targeting advanced prostate cancer with STEAP1 chimeric antigen receptor T cell therapy"

**Supplemental Figure Legends**

**Supplemental Figure 1. Characteristics of STEAP1 expression in lethal mCRPC tissues.** (**A**) Photomicrographs of select mCRPC tissue cores after STEAP1 IHC staining to highlight the plasma membrane staining consistent with staining intensity scores of 0, 1, 2, and 3. Scale bars = 50 μm. (**B**) Plot showing the STEAP1 H-scores of mCRPC tissues cores based on their metastatic site. Dashed line represents a STEAP1 H-score of 30. Plots of STEAP1 H-score and (**C**) AR H-score or (**D**) synaptophysin (SYP) H-score for each mCPC tissue core. For panel **B**, two-way ANOVA with Tukey's multiple comparisons test was used. For panels **C** and **D**, correlation was evaluated using Pearson correlation and respective r square and p-values are indicated.

**Supplemental Figure 2. Validation of the antigen-specific activation and target cell cytolysis of STEAP1-BBζ CAR T cells.** (**A**) Representative flow cytometry plots showing immunophenotyping of CD4 and CD8 T cell products at day 9 of expansion from untransduced control and lentiviral STEAP1 CAR transduction conditions. IFN-γ quantification by ELISA from (**B**) control culture conditions or (**C**) co-cultures of either untransduced T cells or STEAP1-BBζ CAR T cells with the DU145 or DU145 STEAP1 cell lines at a 1:1 ratio at 24 hours. n = 4 replicates per condition. Bars represent SD. (**D**) Relative cell viability of DU145 STEAP1 target cells over time measured by fluorescence live cell imaging upon co-culture with (left) STEAP1-BBζ CAR T cells or (right) untransduced T cells at a 1:1 ratio. n = 4 replicates per condition. Error bars represent SEM. For panels **C** and **D**, unpaired two-tailed Student's t test with Welch's correction was used.

**Supplemental Figure 3. Determination of the STEAP1 ectodomain specificity of STEAP1-BBζ CAR T cells using mouse/human Steap1 chimeras.** (**A**) Immunoblot analysis confirming the lack of human STEAP1 (hSTEAP1) and mouse Steap1 (mSteap1) expression in the DU145 cell line and their respective expression in the lentivirally engineered DU145 hSTEAP1 and DU145 mSteap1 lines. GAPDH is used as a protein loading control. (**B**) IFN-γ quantification by ELISA from (**B**) co-cultures of STEAP1-BBζ CAR T cells with the DU145, DU145 hSTEAP1, and DU145 mSteap1 cell lines at a 1:1: ratio or (**C**) control culture conditions at 24 hours. n = 4 replicates per condition. Bars represent SD. (**D**) Schematic of the mSteap1 protein with extracellular domains (mECDs, blue) highlighted and the mouse/human Steap1 chimeric proteins each with individual replacement of a mECD with the counterpart hSTEAP1 extracellular domain (hECD, red). (**E**) . IFN-γ quantification by ELISA co-cultures of STEAP1-BBζ CAR T cells with each of the DU145 lines engineered to express hSTEAP1, mSteap1, or mouse/human Steap1 chimeric proteins at a 1:1 ratio at 24 hours. n = 4 replicates per condition. Bars represent SD. (**F**) Alignment of the amino acid sequences of the hSTEAP1 ECD2 and mSteap1 ECD2 showing non-conserved residues at positions 198 and 209. Error bars represent SD. For panels **B-D,** two-way ANOVA with Tukey's multiple comparisons was used.

**Supplemental Figure 4. Evaluation of the reactivity of STEAP1-BBζ CAR T cells to STEAP1B isoforms.** (**A**) Alignment of the amino acid sequence of the hSTEAP1 ECD2 to human STEAP1B isoforms 1, 2, and 3 showing complete sequence conservation. (**B**) Plots of TOPCONS consensus predictions of membrane protein topology and reliability scores for hSTEAP1, mSteap1, and the three human STEAP1B isoforms. (**C**) Plots of membrane protein topology predictions for human STEAP1B isoform 1 showing disagreement between different algorithms. (**D**) IFN-γ quantification by ELISA co-cultures of STEAP1-BBζ CAR T cells with each of the DU145 lines engineered to express hSTEAP1, mSteap1, or human STEAP1B isoforms at a 1:1 ratio at 24 hours. n = 4 replicates per condition. Bars represent SD.

**Supplemental Figure 5. Characterization of the expansion, transduction efficiency, and differentiation states of STEAP1-BBζ CAR T cell products.** (**A**) Plots showing the relative expansion of untransduced and STEAP1-BBζ CAR T cell subsets derived from three independent donors over time in culture. (**B**) Table showing the percentage of T cell subsets transduced with STEAP1-BBζ CAR lentivirus that express EGFRt as measured by flow cytometry seven days after transduction. (**C**) Flow cytometry histogram plots showing the expression of the T cell exhaustion markers PD-1 and LAG-3 in PBMCs as well as untransduced and STEAP1-BBζ CAR T cell subsets. T cells treated with PHA-L are used as a positive control for induction of PD-1 and LAG-3 expression. Bar graphs showing the percentages of (**D**) stem cell memory T cells (Tscm: $CD62L^+$ $CD45RA^+$ $CD95^+$ $CXCR3^+$) and (**E**) central memory T cells (Tcm: $CD62L^+$ $CD45RA^-$) in T cell subsets from PBMCs as well as untransduced and STEAP1-BBζ CAR T cell products.

**Supplemental Figure 6. Antitumor activity of STEAP1-BBζ CAR T cell therapy in human 22Rv1 prostate cancer cell line xenograft models.** (**A**) Photomicrograph of CD3 IHC staining of a 22Rv1 subcutaneous tumor 25 days after intratumoral treatment with STEAP1-BBζ CAR T cells. Scale bars = 100 µm. (**B**) Representative photomicrographs of STEAP1 IHC staining of 22Rv1-fLuc subcutaneous tumors after intratumoral treatment with (left) untransduced T cells or (right) STEAP1-BBζ CAR T cells. Scale bars = 50 µm. (**C**) Plot of average weights of NSG mice engrafted wih 22Rv1-fLuc metastatic tumors over time after treatment with either untransduced T cells or STEAP1-BBζ CAR T cells. Bars represent SD. (**D**) Representative low magnification photomicrographs of STEAP1 IHC staining of 22Rv1-fLuc metastatic liver tumors after intravenous treatment with (left) untransduced T cells or (right) STEAP1-BBζ CAR T cells. Scale bars = 1 mm. Higher magnification photomicrographs of (**E**) STEAP1 IHC staining and (**F**) PSMA IHC staining of regions shown in **D**. Scale bars = 50 µm.

**Supplemental Figure 7. Antitumor activity of STEAP1-BBζ CAR T cell therapy in a disseminated human C4-2B prostate cancer cell line xenograft model.** (**A**) Plot of average weights of NSG mice engrafted wih C4-2B-fLuc metastatic tumors over time after treatment with either untransduced T cells or STEAP1-BBζ CAR T cells. Bars represent SD. (**B**) *Ex vivo* BLI of liver and lung tissues from NSG mice engrafted with C4-2B-fLuc metastatic tumors and treated with either (left) untransduced T cells or (right) STEAP1-BBζ CAR T cells. Radiance scale is shown.

**Supplemental Figure 8. Antitumor activity of STEAP1-BBζ CAR T cell therapy in a disseminated human PC3 prostate cancer cell line xenograft model.** (**A**) Plot and table showing absolute quantification of STEAP1 molecules per cell across select human prostate cancer cell lines including PC3 as determined by flow cytometry using external standard beads. (**B**) Relative cell viability of PC3 target cells over time measured by fluorescence live cell imaging upon co-culture with untransduced T cells or STEAP1-BBζ CAR T cells at a 1:1 ratio. n = 4 replicates per condition and bars represent SEM. (**C**) Schematic of tumor challenge experiments for the PC3 disseminated model. (**D**) Serial live bioluminescence imaging (BLI) of NSG mice engrafted with PC3-fLuc metastases and treated with a single intravenous injection of 5 x $10^6$ untransduced T cells or STEAP1-BBζ CAR T cells at a normal CD4/CD8 ratio on day 0. Mice were euthanized due to the onset of severe xenogeneic graft-versus-host (GVHD) in both treatment arms in week 4. Radiance scale is shown. (**E**) Plot showing the quantification of total flux over time from live BLI of each mouse in **F**.

**Supplemental Figure 9. Antitumor activity of STEAP1-mBBζ CAR T cell therapy in a disseminated, non-clonal mouse RM9-hSTEAP1 prostate cancer allograft model in NSG mice.** (**A**) Schematic of tumor challenge experiments for the non-clonal RM9-hSTEAP1 disseminated model. (**B**) Serial live BLI of NSG mice engrafted with RM9-hSTEAP1-fLuc metastases and treated with a single intravenous injection of 5 x $10^6$ untransduced mouse T cells or mouse STEAP1-mBBζ CAR T cells on day 0. Red X denotes deceased mice. Radiance scale is shown. (**C**) Plot showing the quantification of total flux over time from live BLI of each mouse in **B**. (**D**) Kaplan-Meier survival curves of mice in **B** with statistical significance determined by log-rank (Mantel-Cox) test. Plots of weights for each mouse (numbered in **B**) over time in the  mouse (**E**) untransduced T cell treatment group and (**F**) STEAP1-mBBζ CAR T cell treatment group. (**G**) Quantification of $mCD3^+mCD19t^+$ STEAP1-mBBζ CAR T cells by flow cytometry from splenocytes of mice treated with STEAP1-BBζ CAR T cells collected at necropsy. (**H**) Photomicrographs at low and magnification of STEAP1 IHC staining of RM9-hSTEAP1 lung tumors after treatment with (left) mouse untransduced T cells or (right) STEAP1-BBζ CAR T cells. Scale bars = 500 µm.

**Supplemental Figure 10. Antitumor activity of STEAP1-mBBζ CAR T cell therapy in a disseminated, clonal mouse RM9-hSTEAP1 prostate cancer allograft model in NSG mice.** (**A**) Schematic of tumor challenge experiments for the clonal RM9-hSTEAP1 disseminated model. (**B**) Serial live BLI of NSG mice engrafted with RM9-hSTEAP1-fLuc metastases and treated with a single intravenous injection of 5 x $10^6$ untransduced mouse T cells or mouse STEAP1-mBBζ CAR T cells on day 0. Red X denotes deceased mice. Radiance scale is shown. (**C**) Plot showing the quantification of total flux over time from live BLI of each mouse in **B**. (**D**) Kaplan-Meier survival curves of mice in **B** with statistical significance determined by log-rank (Mantel-Cox) test.

**Supplemental Figure 11. Absence of cytokine storm based on serum cytokine analysis before and after mouse STEAP1-mBBζ CAR T cell therapy in hSTEAP1-KI mice.** Plots showing serum cytokine levels (IFN-γ, IL-2, IL-6, and TNF-α) based on ProcartaPlex immunoassays from retroorbital bleeds of hSTEAP1-KI/+ mice bearing RM9-hSTEAP1-fLuc metastases prior to (day 0) or after (day 8) treatment with untransduced mouse T cells or mouse STEAP1-mBBζ CAR T cells. n=3-4 for each condition. Bar represents the mean.

**Supplemental Figure 12. Preserved tissue architecture and absence of increased T cell infiltration in the prostates or adrenal glands of hSTEAP1-KI mice treated with mouse STEAP1-mBBζ CAR T cells.** Representative photomicrographs of hematoxylin & eosin (H&E) and CD3 IHC staining of hSTEAP1-KI/+ prostates from mice treated with (**A**) untransduced T cells and (**B**) STEAP1-mBBζ CAR T cells. Red arrowheads indicate rare $CD3^+$ cells. Scale bars = 50 μm. Representative photomicrographs of H&E and CD3 IHC staining of hSTEAP1-KI/+ adrenal glands from mice treated with (**C**) untransduced T cells and (**D**) STEAP1-mBBζ CAR T cells. Red arrowheads indicate $CD3^+$ cells. Scale bars = 100 μm.

**Supplemental Figure 1**

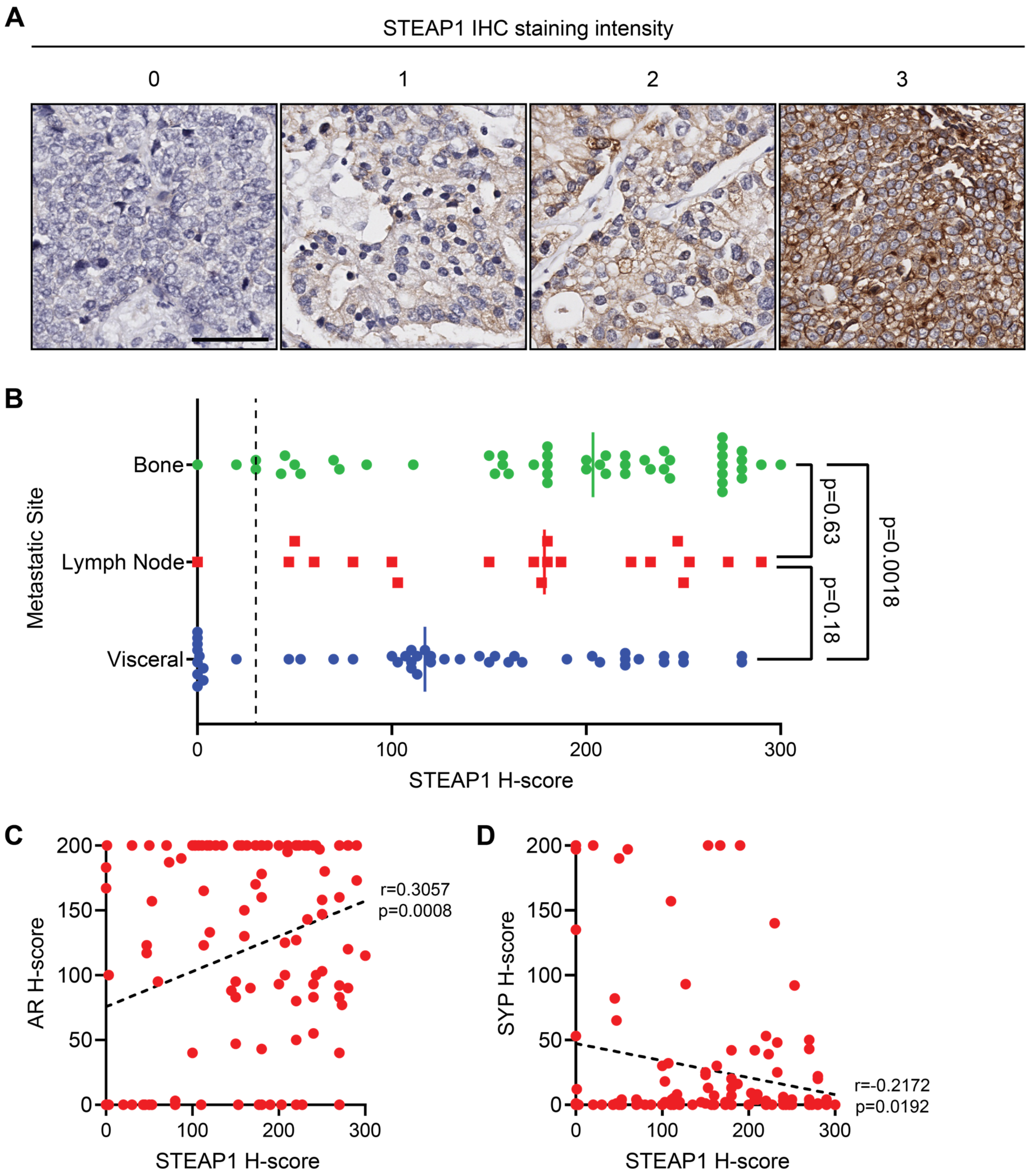

### Supplemental Figure 2

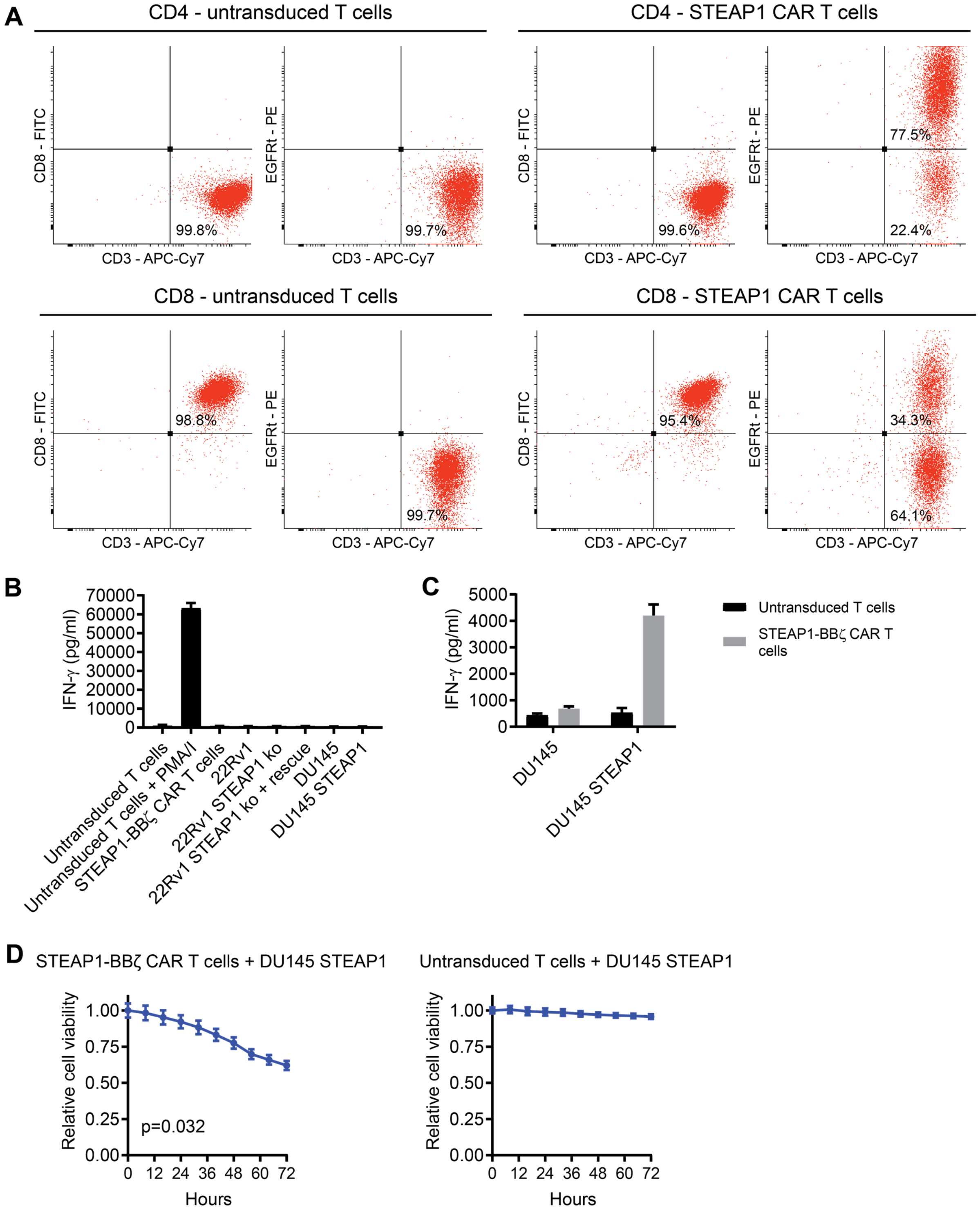

### Supplemental Figure 3

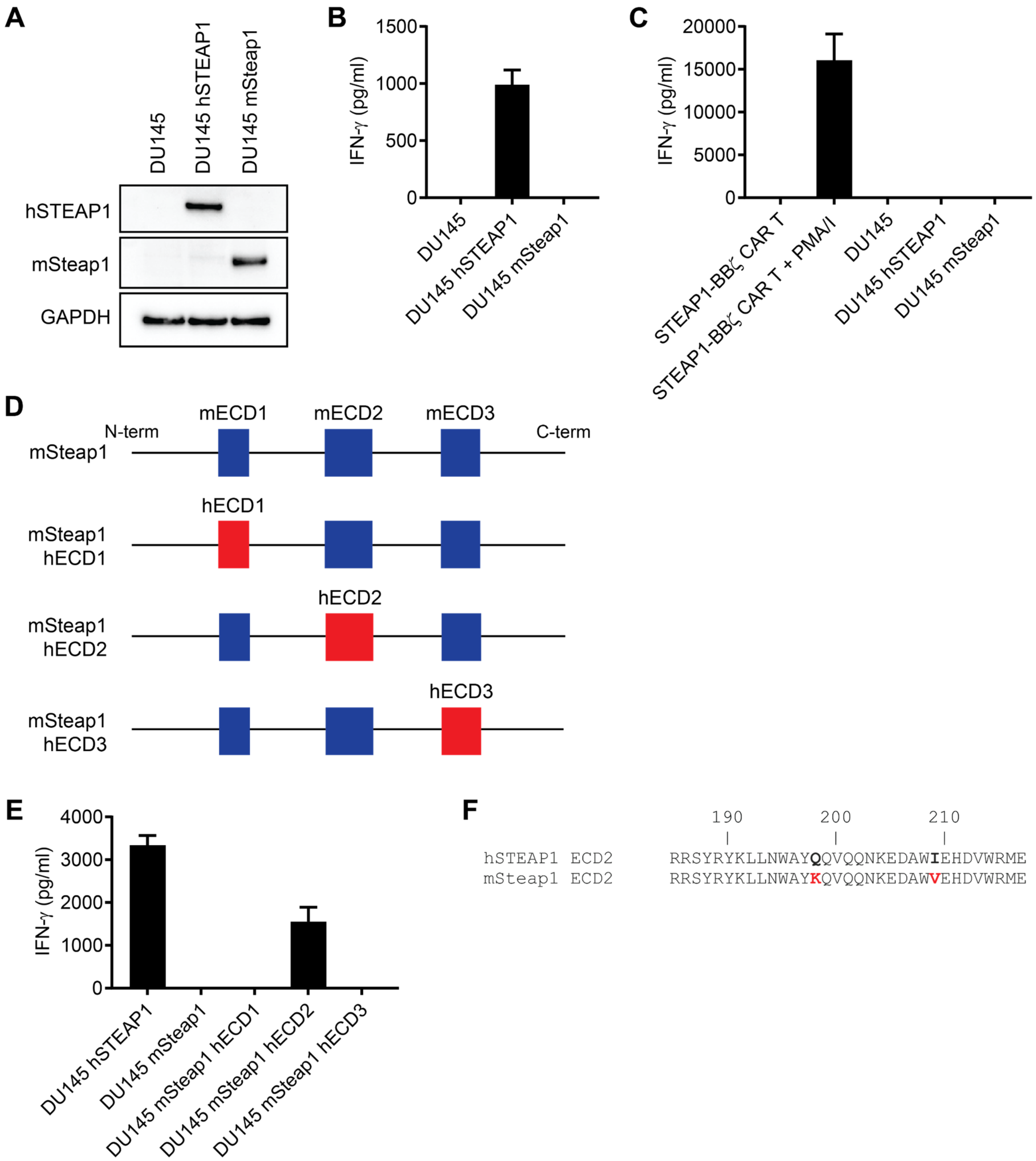

### Supplemental Figure 4

**A**

```
                    190       200       210
                      |         |         |
hSTEAP1 ECD2     RRSYRYKLLNWAYQQVQQNKEDAWIEHDVWRME
STEAP1B iso 1    RRSYRYKLLNWAYQQVQQNKEDAWIEHDVWRME
STEAP1B iso 3    RRSYRYKLLNWAYQQVQQNKEDAWIEHDVWRME

                   170       180       190
                     |         |         |
STEAP1B iso 2    RRSYRYKLLNWAYQQVQQNKEDAWIEHDVWRME
```

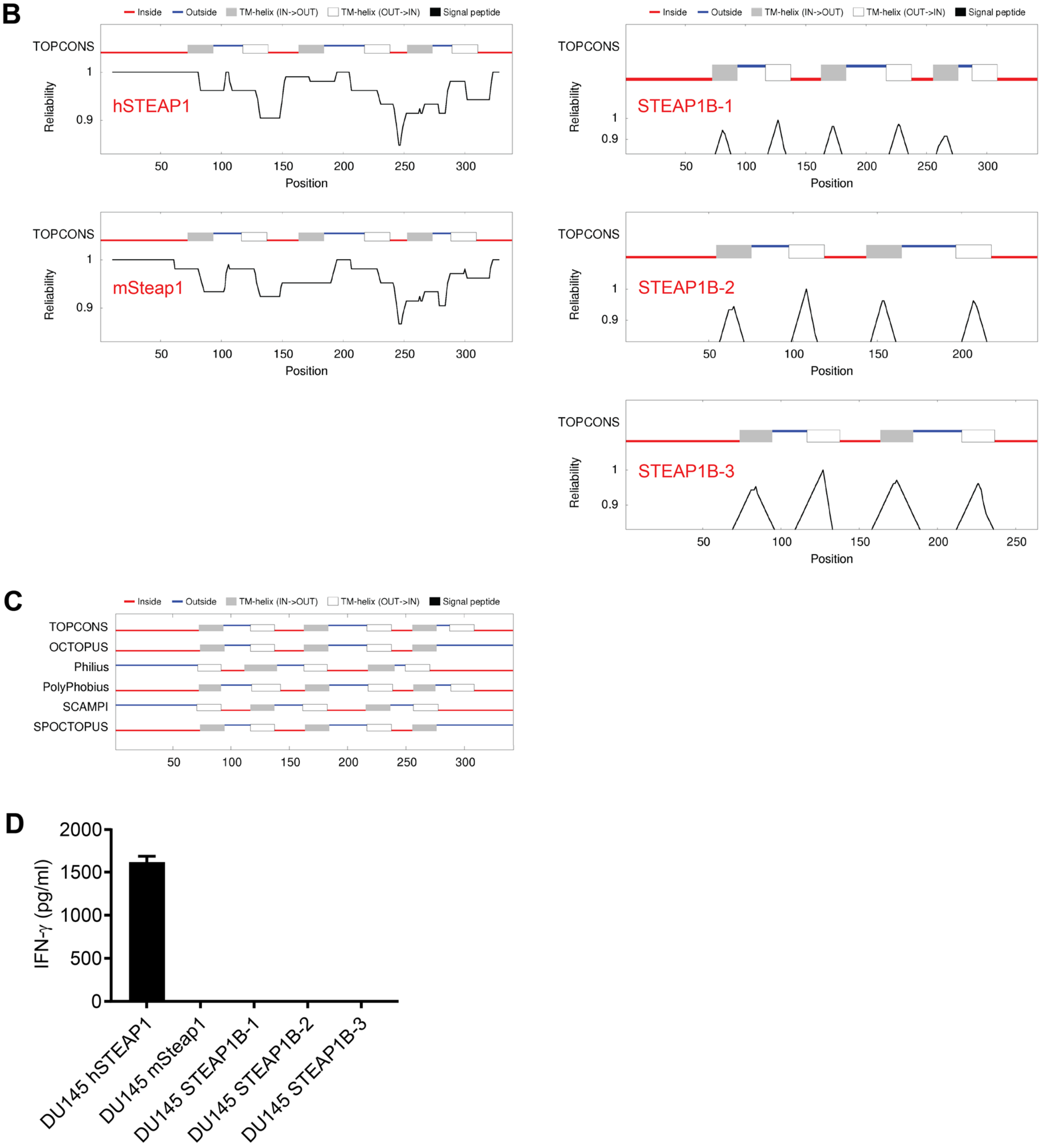

### Supplemental Figure 5

A

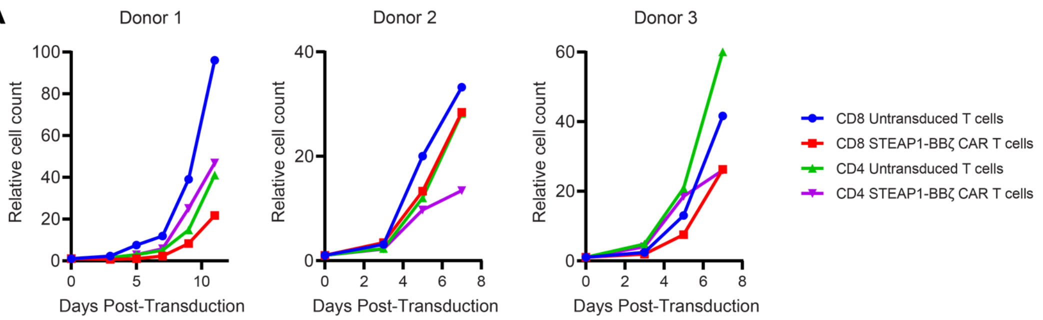


B

| | CD4 EGFRt⁺ | CD8 EGFRt⁺ |
|---|---|---|
| Donor 1 | 60.1% | 26.2% |
| Donor 2 | 74.9% | 24.3% |
| Donor 3 | 69.3% | 54.2% |

C

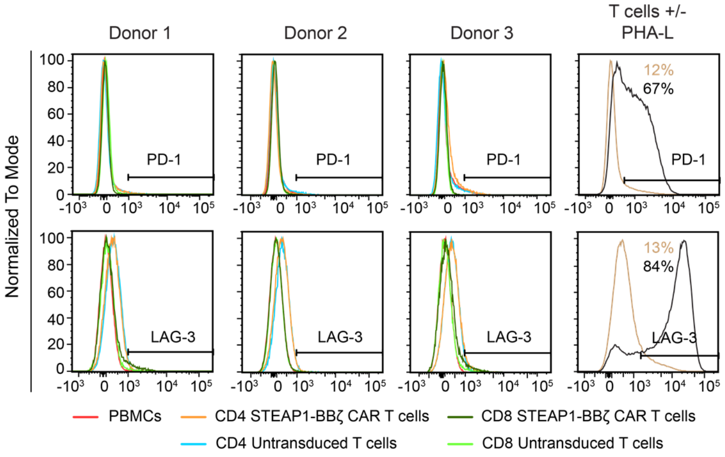


D

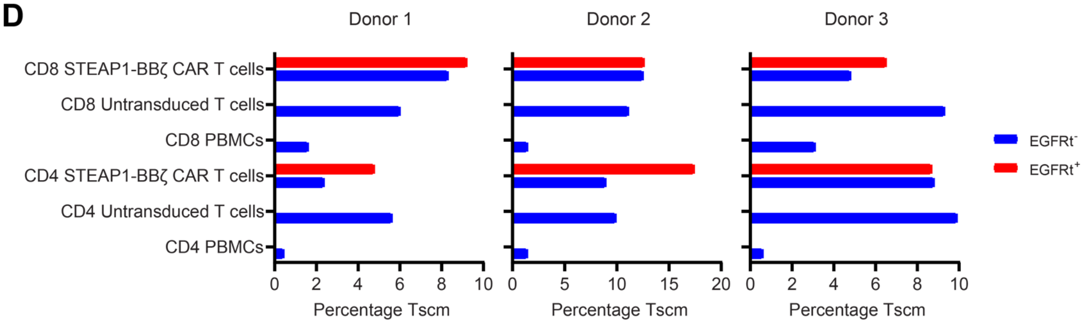


E

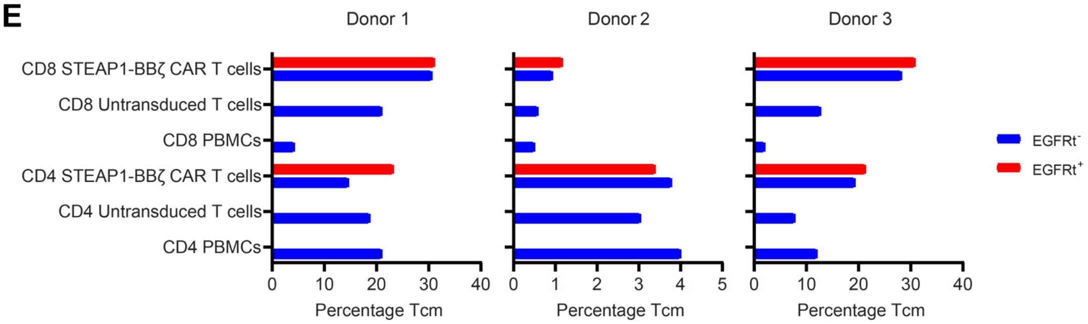

### Supplemental Figure 6

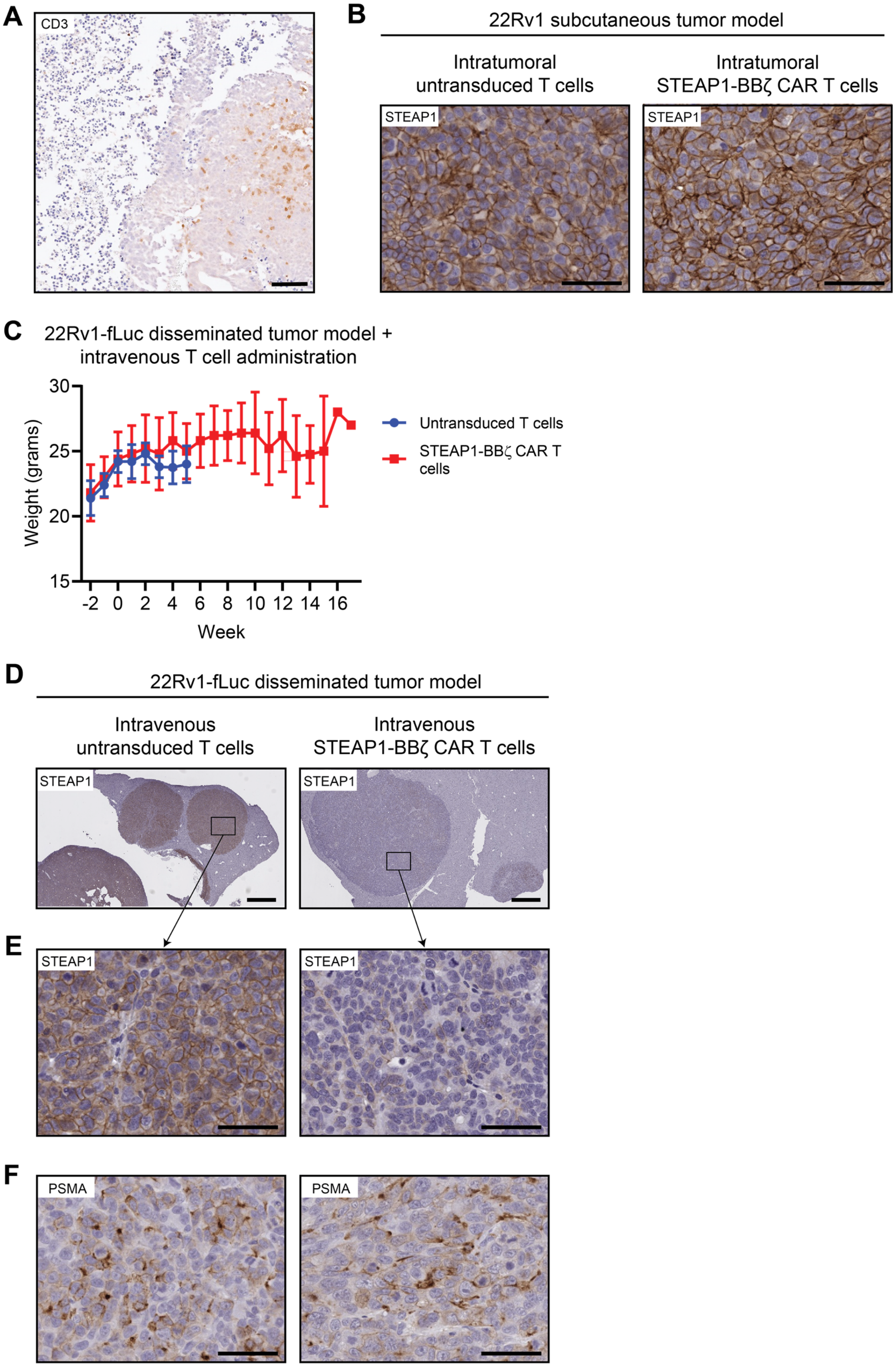

### Supplemental Figure 7

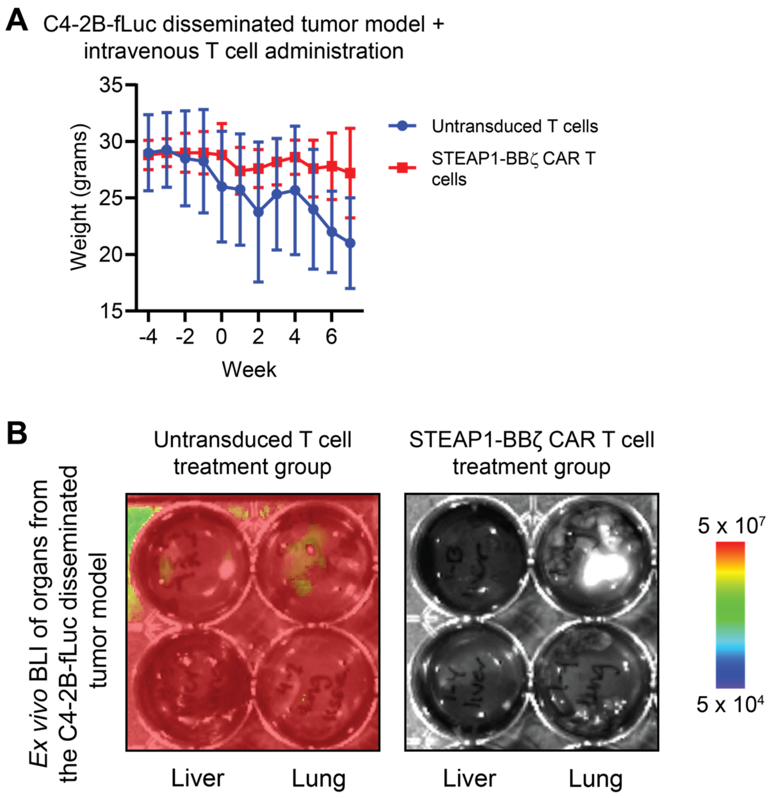

### Supplemental Figure 8

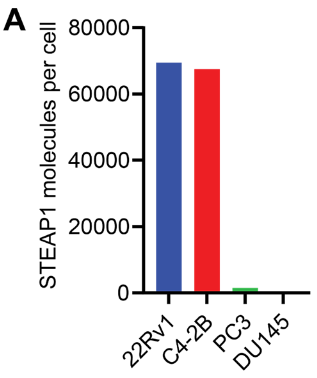


| Cell line | STEAP1 molecules per cell |
|---|---|
| 22Rv1 | 69,475 |
| C4-2B | 67,462 |
| PC3 | 1,491 |
| DU145 | 0 |

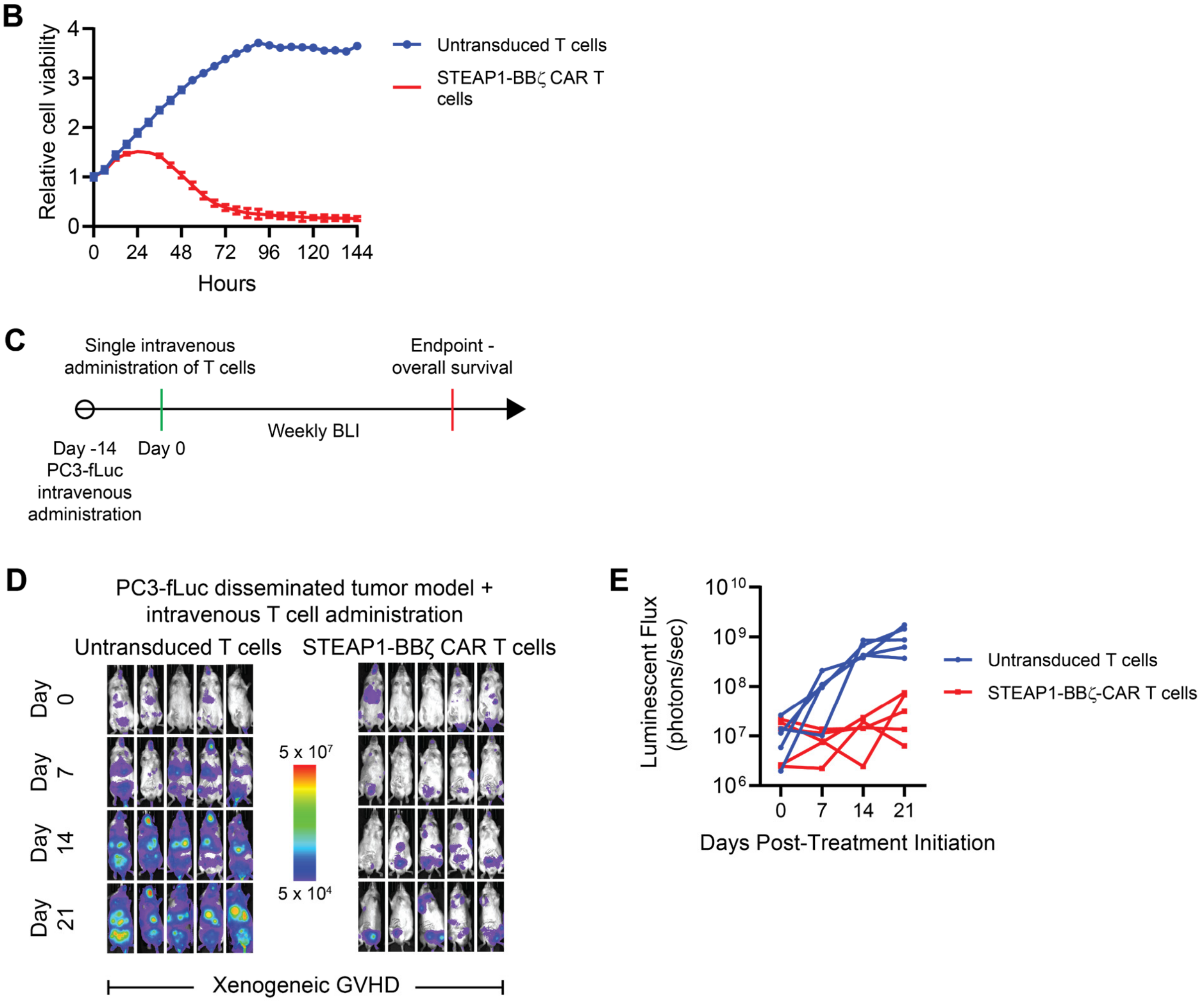

### Supplemental Figure 9

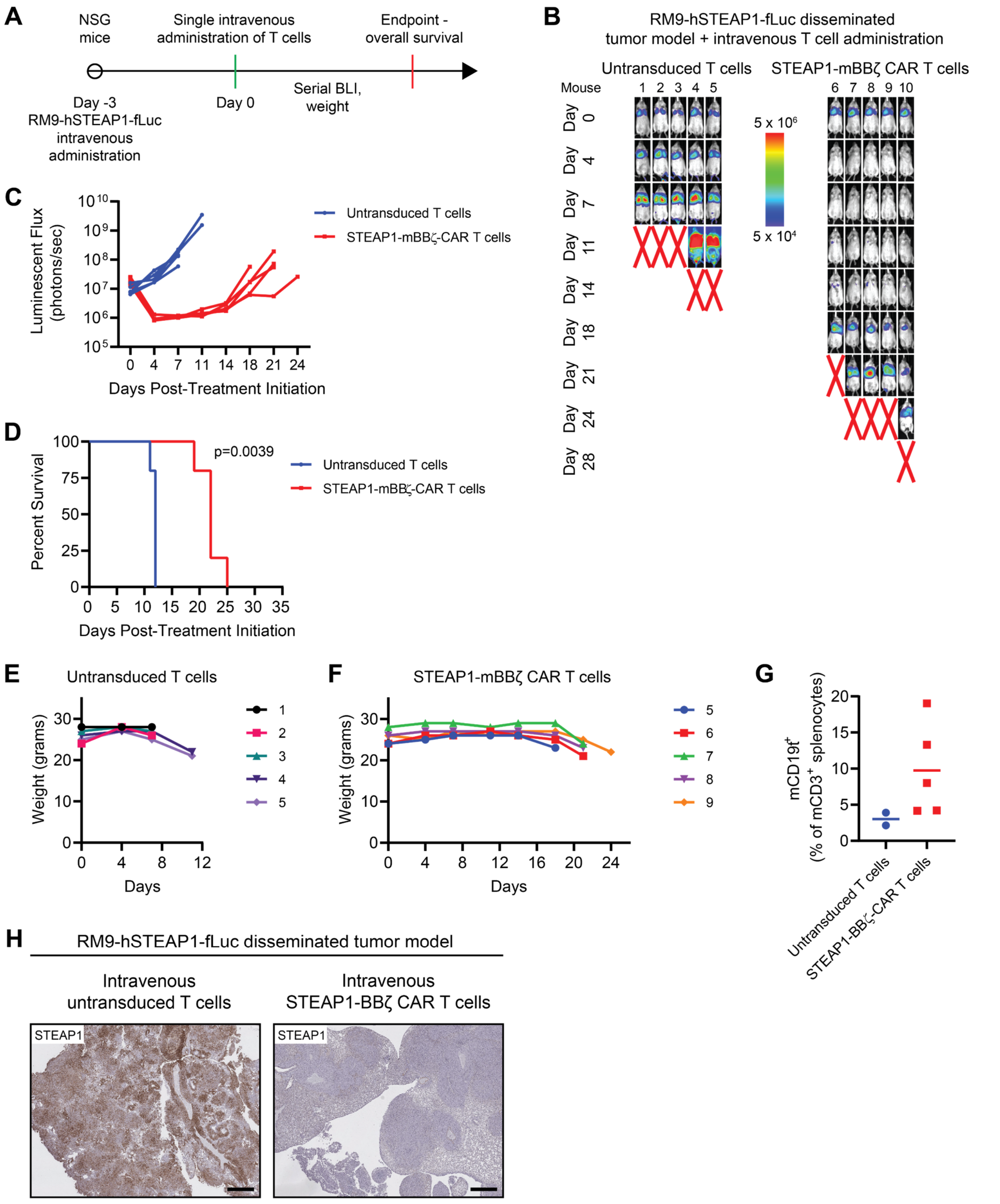

### Supplemental Figure 10

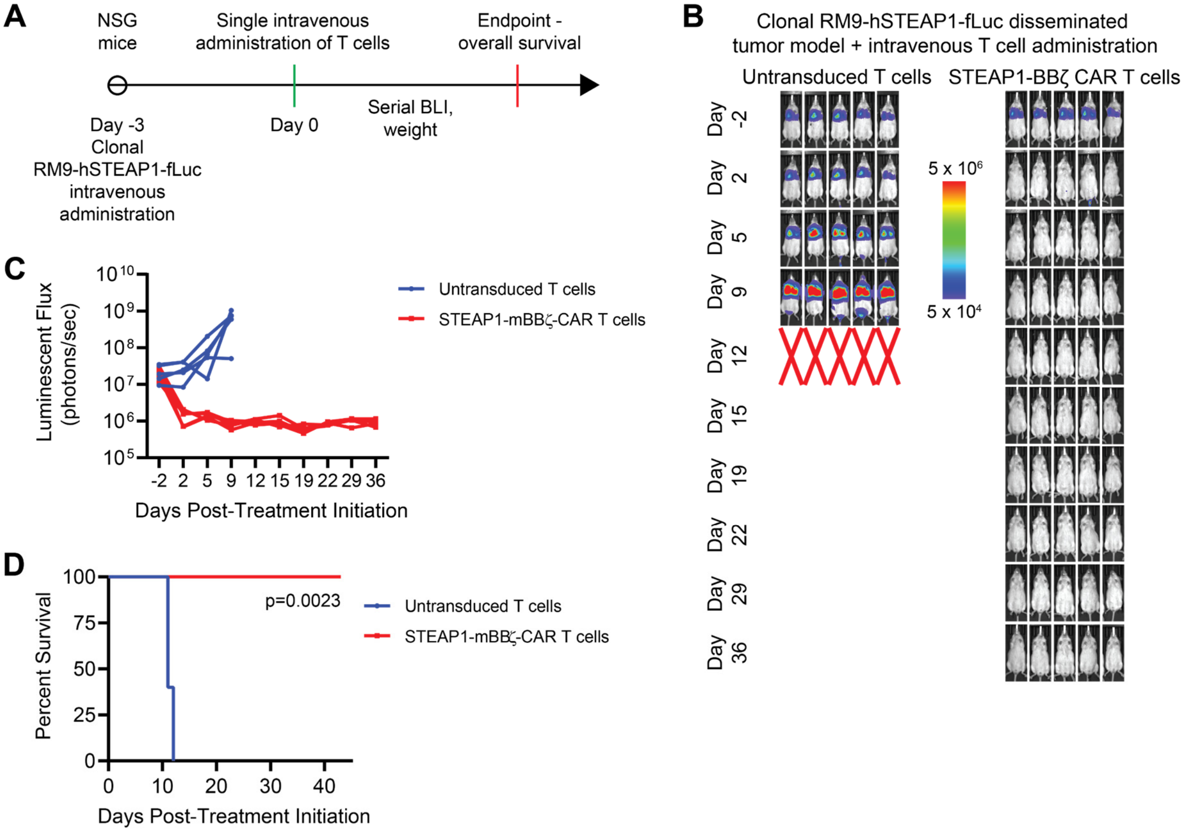

### Supplemental Figure 11

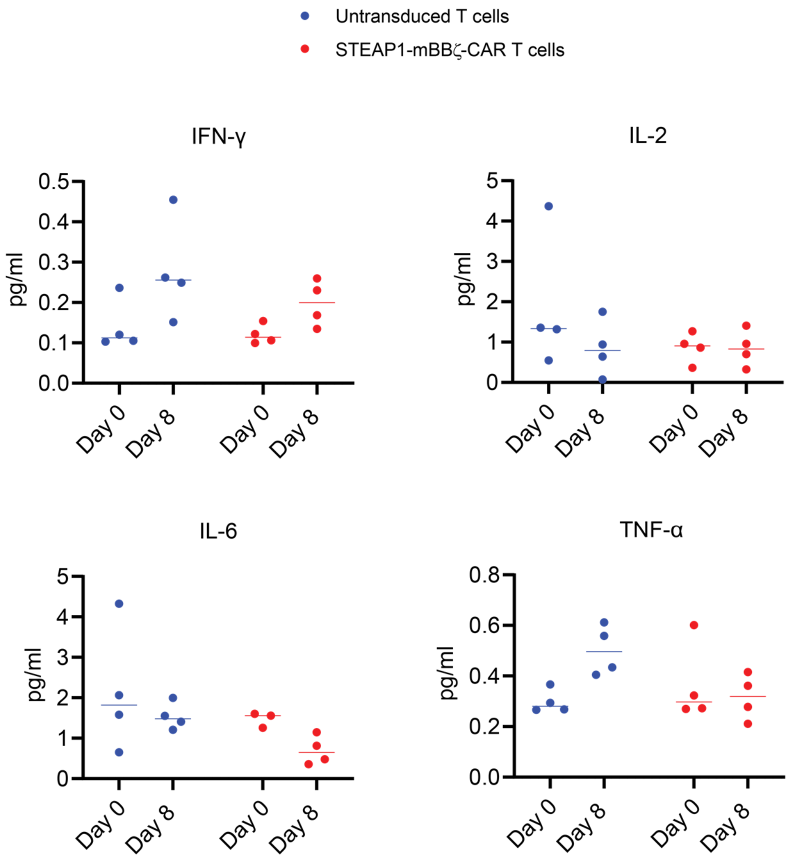

### Supplemental Figure 12

**A** KI/+ prostate from the untransduced T cell treatment group

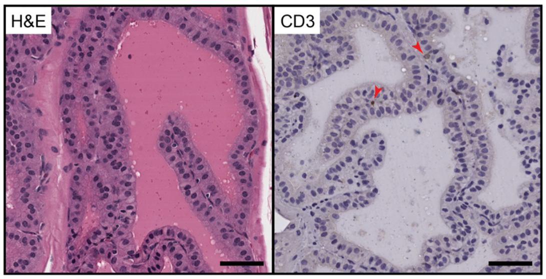


**B** KI/+ prostate from the STEAP1-mBBζ CAR T cell treatment group

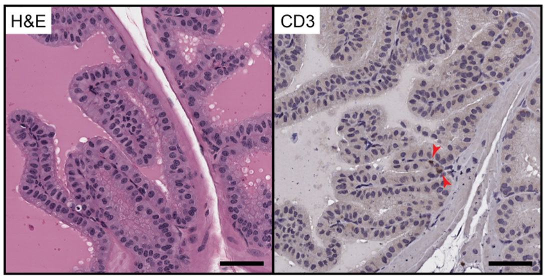


**C** KI/+ adrenal gland from the untransduced T cell treatment group

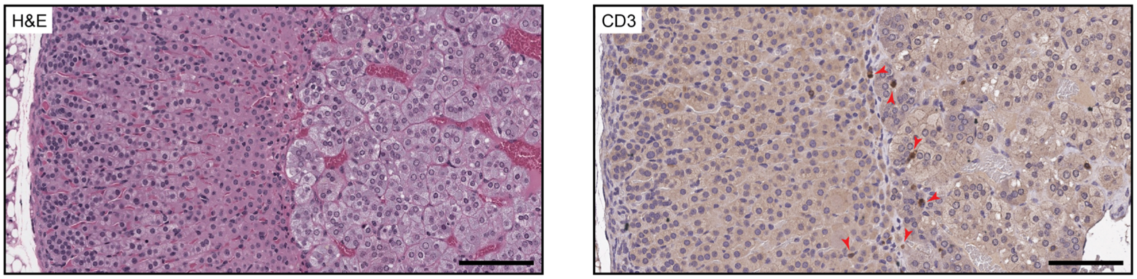


**D** KI/+ adrenal gland from the STEAP1-mBBζ CAR T cell treatment group

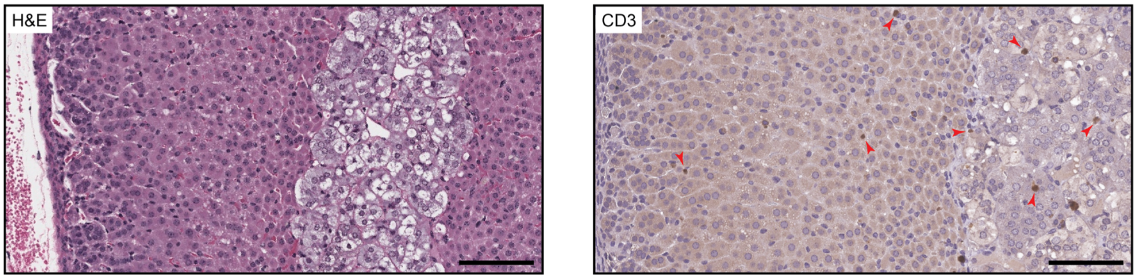
